# Predicting Virus Fitness: Towards a structure-based computational model

**DOI:** 10.1101/2023.05.01.538902

**Authors:** Shivani Thakur, Kasper Planeta Kepp, Rukmankesh Mehra

## Abstract

Predicting the impact of new emerging virus mutations is of major interest in surveillance and for understanding the evolutionary forces of the pathogen. The SARS-CoV-2 surface spike-protein (S-protein) binds to human ACE2 receptors as a critical step in host cell infection. At the same time, S-protein binding to human antibodies neutralizes the virus and prevents interaction with ACE2. Here we combine these two binding properties in a simple virus fitness model, using structure-based computation of all possible mutation effects averaged over 10 ACE2 complexes and 10 antibody complexes of the S-protein (∼3,80,000 computed mutations), and validated the approach against diverse experimental binding/escape data of ACE2 and antibodies. The ACE2-antibody selectivity change caused by mutation (i.e., the differential change in binding to ACE2 vs. immunity-inducing antibodies) is proposed to be a key metric of fitness model, enabling systematic error cancelation when evaluated. In this model, new mutations become fixated if they increase the selective binding to ACE2 relative to circulating antibodies, assuming that both are present in the host in a competitive binding situation. We use this model to categorize viral mutations that may best reach ACE2 before being captured by antibodies. Our model may aid the understanding of variant-specific vaccines and molecular mechanisms of viral evolution in the context of a human host.

## Introduction

The pandemic caused by severe acute respiratory syndrome coronavirus 2 (SARS-CoV-2)^1–3^ has been characterized by the emergence of variants involving mutations in the surface spike glycoprotein (S-protein) that is involved in human host cell entry and is important for immune recognition.^4–6^ The direct health implications of these variants have put the field of protein evolution into a central role, with a public understanding of the importance of mutations leading to more infectious or antibody- or vaccine-resistant variants.^6–11^ Accordingly, there is substantial interest in using protein evolution and protein modelling methods to rationalize and predict these events.

The S-protein enters human host cells by fusion with the cell-surface receptor angiotensin-converting enzyme 2 (ACE2), and the high binding affinity of the S-protein variants towards ACE2 is a prerequisite for infection.^12–14^ However at the same time, the S-protein is presented to the human immune system leading to development of immunity in the population after infection or after vaccination, since this presentation is the design rationale behind many vaccines.^15, 16^ These two binding processes can be viewed as opposed, with one favoring infection and the other limiting it, and thus, the fitness of the virus is linked to both these binding events, as explored further in this study.

Evolution occurs by changing mutations in a folded protein structure,^17–21^ and thus it is important to provide the structural context to the mutations in order to predict and rationalize their impacts. Fortunately, the few years before the pandemic witnessed major technical breakthroughs in the field of cryo-electron microscopy applied to solving the 3-dimensional structures of macromolecules.^22–25^ This development made it possible during the pandemic for research groups to publish hundreds of structures of the S-protein solved for all relevant conformational states either by itself or in complex with ACE2 or a large variety of antibodies, typically at resolutions of 2-4 Å that establish well the polypeptide backbone structures.^26^ This means that structure-based evolution models of the SARS-CoV-2 S-protein are in principle feasible.

The present work develops a new method by using ensembles of these experimental structures and computational models estimating binding affinities to evaluate mutation effects in a model of virus fitness that includes both ACE2 and antibody binding as a competitive binding phenomenon. We propose that the host-virion interaction can be viewed as a situation where the virion seeks to enter the cell via binding its S-protein to ACE2 before the S-protein is bound by circulating antibodies. In this model, the virus fitness becomes directly proportional to the binding affinity towards ACE2 minus the binding affinity towards a representative cocktail of antibodies, and the mutation effect on fitness (the selection coefficient) becomes proportional to the change in this difference relative to the wild-type or reference strain.

One of the advantages of this “selectivity model” is that it compares differences of differences (differences in binding differences of mutant and wild type to ACE2 and antibodies). This protocol removes systematic errors that otherwise always exist in the experimental assays and especially in the computer models, from bias towards some amino acid mutation types^27, 28^ and reliance on a folded wild-type structure for estimating the mutation effects.^27–35^ Our results correlated well with diverse experimental binding/escape datasets for ACE2 and antibodies. Overall, our work demonstrates a simple, intuitively appealing, and computable model to estimate the fitness function of the virus.

## Methods

### Strategy and hypothesis

ACE2 and antibodies (AB) both bind to the S-protein. ACE2 binding induces conformational changes in the S-protein, mainly in the receptor binding domain (RBD). AB binding may also be associated with conformational changes and the binding sites of different ABs may vary, although many bind to the RBD.

We hypothesize that the virus particle, in order to be successful, needs to bind with its S-protein to ACE2 *before* being bound to an antibody, and we assume that the ACE2 receptors and antibodies are available in a competitive binding equilibrium (the relative amount of them may differ, but this is supposed to be the same condition for all variants when evaluating the fitness effects, and if so, does not affect the model). Under these conditions, the selectivity (the differential binding affinity to ACE2 vs. the ensemble of circulating antibodies) defines the fraction of virions that infects cells before they are neutralized by antibodies of the immune system. We also propose that this selectivity defines a fitness function of the virus as it accounts both for optimized binding to ACE2 and antigenic drift (selection pressure to weaken binding to antibodies). Combining the two binding affinities directly seems important because many mutations may strengthen binding to many proteins including both ACE2 and antibodies, but it is the *differential* binding effect that we envision is important.

Since cryo-EM structures and computer models are available to compute the binding affinity change caused by mutation, we can compute these for an ensemble of structures to get more robust results given the structural heterogeneity, for both ACE2 and an ensemble of antibodies, and subtract these binding effects of mutation, to yield a fitness function F:

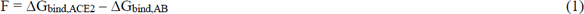

Where ΔG_bind,AB_ is the binding free energy of the S-protein of the variant to the weighted ensemble of circulating antibodies, and ΔG_bind,ACE2_ is the binding affinity to ACE2 for the same variant. The selection coefficients then become proportional to the change in fitness:

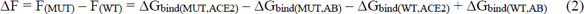

Or written in a shorter form,

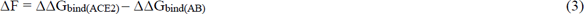

A major advantage of this model is not only that it combines ACE2 and antibody binding directly in a similar treatment, but also that this reduces systematic errors on the final metric *s* (the selectivity coefficient, which is the difference in fitness ΔF of the mutant and wild type or reference strain). Since any denominator of the selection coefficient is a constant for the purpose of the present work (using the same reference strain, the Wuhan variant), we have set it to 1 in the following to reduce the computation complexity, and thus only consider ΔF. The values thus need to be scaled with a constant if estimating true selection coefficients, which is beyond the scope of the present work.

### Protein structural datasets

To develop the model described above, we need to use an ensemble of structures, since using one structure for each of the two cases is insufficient due to the variation in ABs requiring an “ensemble” or cocktail of these to be studied, and also, even for the same chemical system, due to coordinate heterogeneity for published structures necessitating an ensemble treatment.^36^ We thus report below average binding effects calculated from ensembles of ten ACE2 and ten AB structures to reduce the impact of these variations.

Specifically, we used two structural datasets: 10 AB-(**Figure S1** and **Table S1**) and 10 ACE2-S-protein (**Table S2)** complexes were selected from Protein Data Bank (PDB). The criteria for the selection include: **(i)** RBD should preferably be in down conformation in AB complexes whereas up in ACE2. **(ii)** For the S-protein-AB complexes, AB should be bound to RBD. **(iii)** The S-protein should be represented by a nearly full-length structure (N > 900, N: number of residues in each chain, **Table S3**). **(iv)** Structures should be of good resolution. **(v)** In addition, all ABs are human monoclonal ABs obtained from SAR-CoV (S309) and SARS-CoV2 infected patients.^37–38^ These criteria ensured consistency in the two datasets (10 ACE2 and 10 AB), binding to the RBD and having similar-length structures to reduce noise and maintain a good level of uniformity for comparing S-protein-ACE2 and -AB complexes with RBD binding regions (**Table S4b**).

**Figure S1** shows how the antibodies bind to the RBD (**Supplementary Section 1**). These structures include 6WPS by Pinto et al.^37^, 6XEY by Liu et al.^39^, 7JV6 by Piccoli et al.^40^, 7K8S and 7K90 by Barnes et al.^41^, 7ND7 by Dejnirattisai et al.^42^, 7LS9 by Cerutti et al.^43^, 7K43 by Tortorici et al.^44^, 7M6E by Scheid et al.^45^ and 7L56 by Rapp et al.^38^ In six structures (**Figure S1a-S1f),** one antibody unit (one heavy and one light chain) is bound to each RBD, whereas in the remaining four structures (**Figure S1g-S1j**), two antibody units interact with each RBD (**Table S1**). We included these two subsets to ensure that we cover all possible AB binding regions on RBD and broad S-protein conformations, since the ensemble of these conformations might form an active AB interacting state. Residue-specific details regarding the binding interface between antibodies and S-protein are shown in **Tables S5-S6**.

The PDB codes of ten S-protein-ACE2 complexes include 7A98 by Benton et al.^46^, 7CT5 by Guo et al.^47^, 7DX8 and 7DX9^48^ by Yan et al.^48^, 7KJ3 and 7KJ4 by Xiao et al.^49^, 7KMS, and 7KMZ, 7KNH, and 7KNI by Zhou et al.^50^ Out of these, six structures have all RBD in the up-conformation (7A98, 7CT5, 7DX9, 7KJ4, 7KMS, 7KNI) and four with two up and one down conformation (7DX8, 7KJ3, 7KMZ, 7KNH). Five structures are three-ACE2 bound, and the other five are two-ACE2 bound S-protein complexes (**Table S2**). By considering multiple structures of these types, we account for such variability in our estimates. In summary, the main results reported below are, unless otherwise noted, based on computed numbers averaged over all the 10 structures of the AB or ACE2 S-protein complexes, respectively.

### Comparative estimates of binding affinity changes in AB vs. ACE2 datasets

In total, we computed the change in binding affinity upon mutation (ΔΔG_bind_ in kcal/mol) in 20 structures (ten S-protein-AB and ten S-protein-ACE2 complexes, three chains in each structure) using the BeAtMuSiC^51^ program, which has yielded promising accuracy in independent benchmarks.^52, 53^ Mutations were introduced simultaneously in all identical chains, and then used to compute the ΔΔG_bind_ = ΔG_bind (MUT)_ − ΔG_bind (WT)_, i.e., the change in binding free energy due to mutation. The negative value indicates an increase in binding affinity and *vice-versa*. Each structure had approximately ∼1000 residues, giving a total of ∼19,000 mutations (19 mutations × 1000) per structure, leading to a total of ∼3,80,000 computational mutation calculations in this work.

### Investigating groups of mutations

Computational mutation models suffer from some limitations partly due to the biased training datasets, the algorithm’s accuracy, the ignorance of structural heterogeneity, and the assumption that mutant properties can be derived from a static wild-type structure.^27, 28, 31, 54^ We have previously shown that the average mutation effect of an ensemble of structures of a group of mutations compared to a control group of mutations is necessary to reduce systematic errors in computational models, much in the same way as in randomized controlled trials but applied in the context of computational (bio)chemistry.^55, 56^ In this way we can determine if a group of mutations in e.g., a variant, have properties significantly different from those of a reference strain, with much higher precision and accuracy than standard absolute estimates.

To this end, the S-protein mutation dataset for each S-protein-AB and -ACE2 complexes was divided into nine subsets (**Table S4**): **(*a*)** “Full length” – all possible mutations in S-protein chain with N residues, i.e., N × 19 mutations; **(*b*)** “Interface” – including mutations in the binding interface residues of the S-protein (listed in **Table S5**). BeAtMuSiC identifies interface residues based on the condition that the difference between a residue’s solvent accessibility in the complex and apo-protein is at least 5%^51^; **(*c*)** “All natural mutations studied” – mutations observed in the wild in different variants; **(*d*)** “Saturation mutagenesis at the natural mutation site” – all possible mutations at the natural mutation sites; **(*e*)** “Interface natural mutations” – those natural mutations found in the S-protein’s interface residues; **(*f*)** “Gamma mutations” – mutations found in the gamma (P.1) variant; **(*g*)** “Delta mutations” – mutations found in the delta (B.1.617.2) variant; **(*h*)** “Omicron mutations” – mutations found in omicron (B.1.1.529) variant; and **(*i*)** “RBD escape mutations” – selected RBD mutations showing strong escape from some antibodies (**Table S7**). The datasets are described in additional detail in **Supplementary Section 2**.

### Statistics

Statistical analyses were performed in the following ways: **(i)** Linear relation of average ΔΔG_bind_ per residue between AB-(ΔΔG_bind(10 AB)_) and ACE2-bound (ΔΔG_bind(10 ACE2)_) S-protein structures. The comparison was made for all possible S-protein mutations in the positions common to both sets of structures (10 AB and 10 ACE2 structures; 20197 data points). The correlation was also checked for the RBD mutations (3819 points), all natural mutations studied (70 points), and RBD natural mutations (25 points) (Figure 1). **(ii)** Linear regression of average ΔΔG_bind(10 ACE2)_ versus ΔΔG_bind(each AB)_ (**Figure S2**), and average ΔΔG_bind(10 AB)_ versus ΔΔG_bind(each ACE2)_ (**Figure S3**). For linear regressions, p-values were calculated. These analyses showed the pattern of ΔΔG_bind_ and how results depended on the choice of structures.

**Figure 1.**
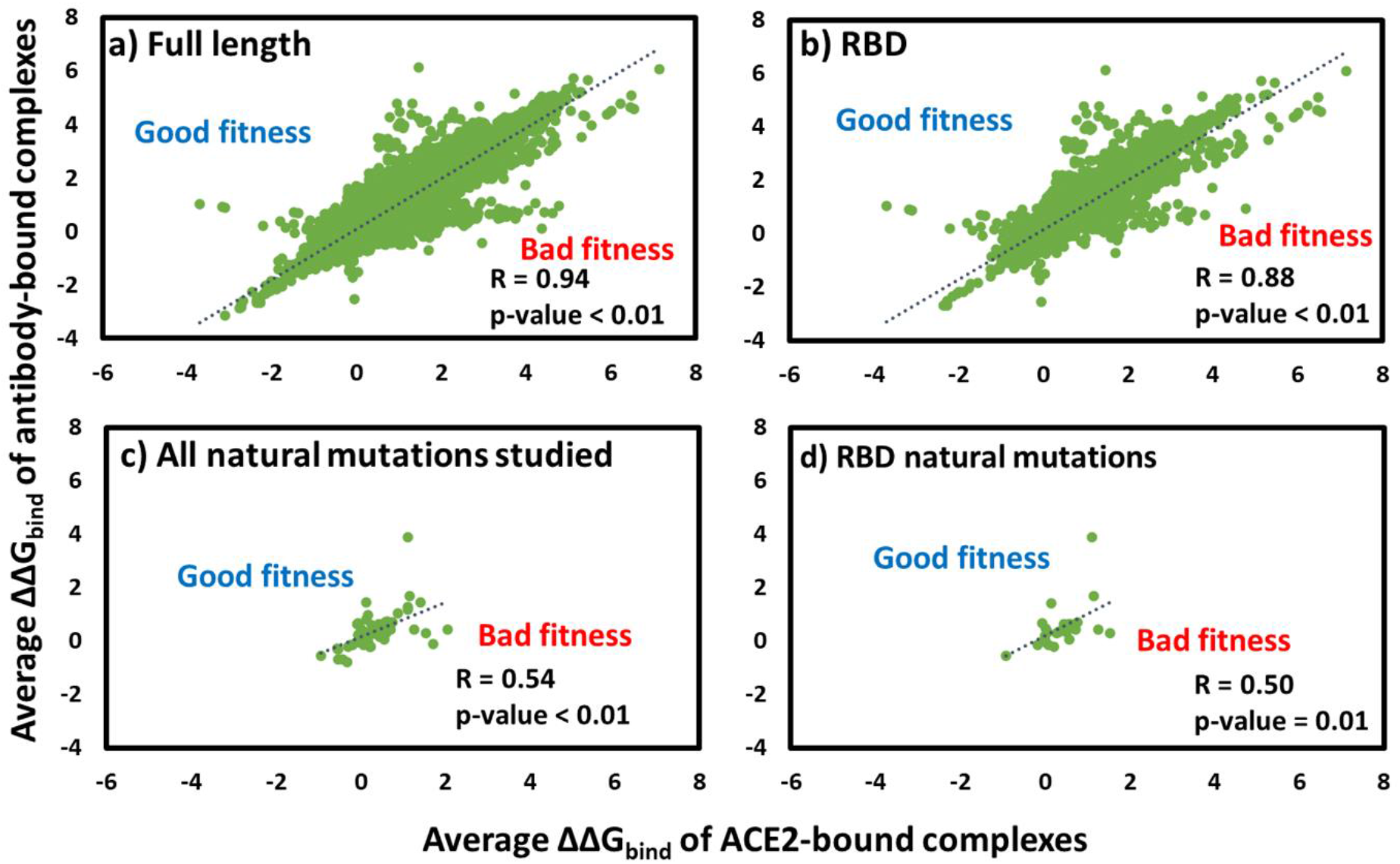
Linear relation between ΔΔG_bind_ values of S-protein mutations bound to AB and ACE2. **(a)** Full length mutations. **(b)** Mutation in RBD residues. **(c)** All natural mutations studied. **(d)** RBD natural mutations.

In addition, we determined significant differences in the structure-averaged binding effects of the mutation groups to AB and ACE2 structures, applied to comparisons of the nine mutation groups defined above (**Table S8**). A Student’s t-test comparison for the same mean binding effect was evaluated for the various combinations of mutation categories (36 combinations) for the AB-/ACE2-bound structures (**Tables S9-S10**).

### Calculating the change in fitness function

As described above, we hypothesized that natural selection of SARS-CoV-2 variants balances antigenic drift with enhanced or maintained binding to ACE2. To model this, we computed the difference in change in affinity of ACE2-bound and AB-bound complexes due to specific mutations, i.e., the differential impact on affinity towards the two targets caused by a single-point substitution. In this way, the selection coefficient (without normalization to the wild-type effect) is proportional to the change in fitness (ΔF) for each mutation, as described by equations (1-3) defined above. We used three approaches to calculate the change in fitness (ΔF) for all natural mutations studied: (i) ΔF = Average ΔΔG_bind(MUT, 10 ACE2)_ – ΔΔG_bind(MUT, each AB)_, (ii) ΔF = ΔΔG_bind(MUT, each ACE2)_ – Average ΔΔG_bind(MUT, 10 AB)_, (iii) Average ΔF = Average ΔΔG_bind(MUT, 10 ACE2)_ – Average ΔΔG_bind(MUT, 10 AB)_.

### Validation of the approach using experimental data

To validate our findings, we compared experimental data in three ways: Firstly, experimental ACE2 binding data from Starr et al. ^57^ (**dataset 0**) as shown in **Table S11** was correlated with computed average ΔΔG_bind_ values of ACE2-bound (3283 data points) complexes. The bin method was used to see the change in correlation for a range of experimental binding, as reported earlier.^58, 59^ The observed ACE2 binding data was only available for RBD mutations, and in the data, negative values indicate a decrease in ACE2 binding and *vice-versa*.

In the second validation approach, we compared experimental AB escape fraction data with the computed average ΔΔG_bind_ values of mutations. We took experimental data of SARS-CoV2 neutralizing ABs from two data sets (**datasets 1 and 2; Table S11**). **Dataset 1** of 1924 data points was the average mutation escape of ten ABs, of which two were also present in our studied structures (C002 in 7K8S and C144 in 7K90 structure). **Dataset 2** comprised 2346 data points, which was also the average mutation escape of ten ABs and involve ABs different from those present in our studied AB structures. In addition, we also correlated the computed binding of individual antibodies C002 (1950 data points) and C144 (1949 data points) with the experimental mutation escape according to **dataset 1**.

Lastly, the experimental differences in ACE2 binding (**dataset 0**) and AB escape (**datasets 1** and **2**) were correlated with computed binding differences (change in fitness, ΔF, equation 3) for RBD mutations. Experimental average AB escape values (**datasets 1** and **2; Table S11**) were subtracted from experimental ACE2 binding values (**dataset 0**), and further, these experimental difference (ACE2 – AB) for each mutation was correlated with the computed binding difference (computed change in fitness **=** Average ΔΔG_bind(MUT, 10 ACE2)_ - Average ΔΔG_bind(MUT, 10 AB)_). The data points involved in two correlations are 1847 (**dataset 1**) and 2245 (**dataset 2**) respectively.

For all comparisons, mutation escape fractions were converted to −log values. The experimental data (**datasets 0, 1** and **2**) were highly squeezed and therefore, the bin method was used with 0.25 bins to see the change in correlation for a range of experimental binding/escape, as reported earlier.^58, 59^

### Estimating epistatic effects

Epistatic effects were estimated using FoldX^60^, by calculating the change in folding stability upon mutation (ΔΔG) after introducing all mutations simultaneously for a given variant, i.e., for the combined effect of mutations allowing them to interact, for gamma 1, gamma 2, delta, and omicron, and additionally selected AB escape mutations (**Table S12)**. The average ΔΔG values were calculated using the ΔΔG values of eight structures of the apo S-protein (average ΔΔG_(apo)_), AB-bound protein (average ΔΔG_(AB)_), and the ACE2-bound S-protein (average ΔΔG_(ACE2)_) for each S-protein variant. The apo S-protein dataset include 6VXX^61^, 6X6P^62^, 6X79^63^, 6XF5^64^, 6Z97^65^, 7CAB^66^, 7DDD^67^ and 7DF3^68^. The S-protein-AB (holo, AB) structures include 6WPS^37^, 6XEY^39^, 7JV6^40^, 7K8S^41^, 7K43^44^, 7K90^41^, 7M6E^45^ and 7ND7^42^. Whereas the S-protein-ACE2 (holo, ACE2) dataset comprises 7DX8^48^, 7DX9^48^, 7KJ3^49^, 7KJ4^49^, 7KMS^50^, 7KMZ^50^, 7KNH^50^ and 7KNI^50^ complexes.

## Results and discussion

### Computed affinities in S-protein-antibodies versus -ACE2 structures

Figure 1 summarizes the impact of all possible mutations averaged over the ten used structures on S-protein binding to ACE2 and the ten studied antibodies. We have previously shown that computed ACE2 binding affinity changes correlate surprisingly well with the experimental data from Bloom et al. once the data sets were curated.^58^ Figure 1 shows that the correlation coefficients illustrate well the expectation that in general, mutations that tend to cause decreased S-protein affinity to one type of proteins (ACE2) also tend to reduce binding to the other type of proteins (antibodies), i.e., the mutation effects have some generic protein-protein interaction effects, related to e.g., hydrophobic packing. Results in **Figures S2-S3** also confirm this, which shows good correlation for the comparison of each AB structure with the average of 10 ACE2 (**Figure S2)**, and *vice-versa* (**Figure S3**).

From an evolution point of view and related to the hypothesis of this work, outliers deviating from the regression line in Figure 1 are of interest as having a potential fitness effect in a competitive binding model, as the binding is not similar for the two protein targets. To understand these outliers better, we separately looked at all mutations together (Figure 1a), all mutations in the RBD (Figure 1b), all studied naturally occurring mutations (Figure 1c) and natural mutations in the RBD (Figure 1d). From Figure 1c, we found Y145D, W152C, E484A, F490S, and Y505H as especially showing differential binding (>1 kcal/mol) between AB and ACE2 structures. Interestingly, most of these (E484A, F490S, and Y505H) are located on the RBD (Figure 1d). Natural mutations Y145D, W152C and Y505H are estimated to have higher binding affinities for antibodies, while E484A and F490S have higher computed binding affinities for ACE2. The F490S mutant is reported to escape several monoclonal antibodies.^69^

### Natural mutations in S-protein in antibody and ACE2 complexes

To clarify the fitness effects of Figure 1 on a residue basis, the ΔΔG_bind_ values for the 70 studied natural S-protein mutations binding to ACE2 and the studied antibodies are summarized in the heatmap in Figure 2 (Note: Out of 79 natural mutations in the dataset, only 70 were present in the experimental S-protein structures and thus computable). The values for each of the 10 structures and the average are shown in the heatmap. The green color represents increased affinity, whereas red shows decreased affinity. Neutral effects are represented in yellow. White spaces indicate missing mutation sites in the respective structures. The heatmap shows differential average affinity changes, i.e., effect on AB/ACE2 selectivity. For mutations with only a few coordinates available (mostly white patches in Figure 2), no significant conclusion could be drawn due to variation in the ΔΔG_bind_ values, which shows the importance of accounting for structural heterogeneity by averaging, as discussed before.^36^

**Figure 2.**
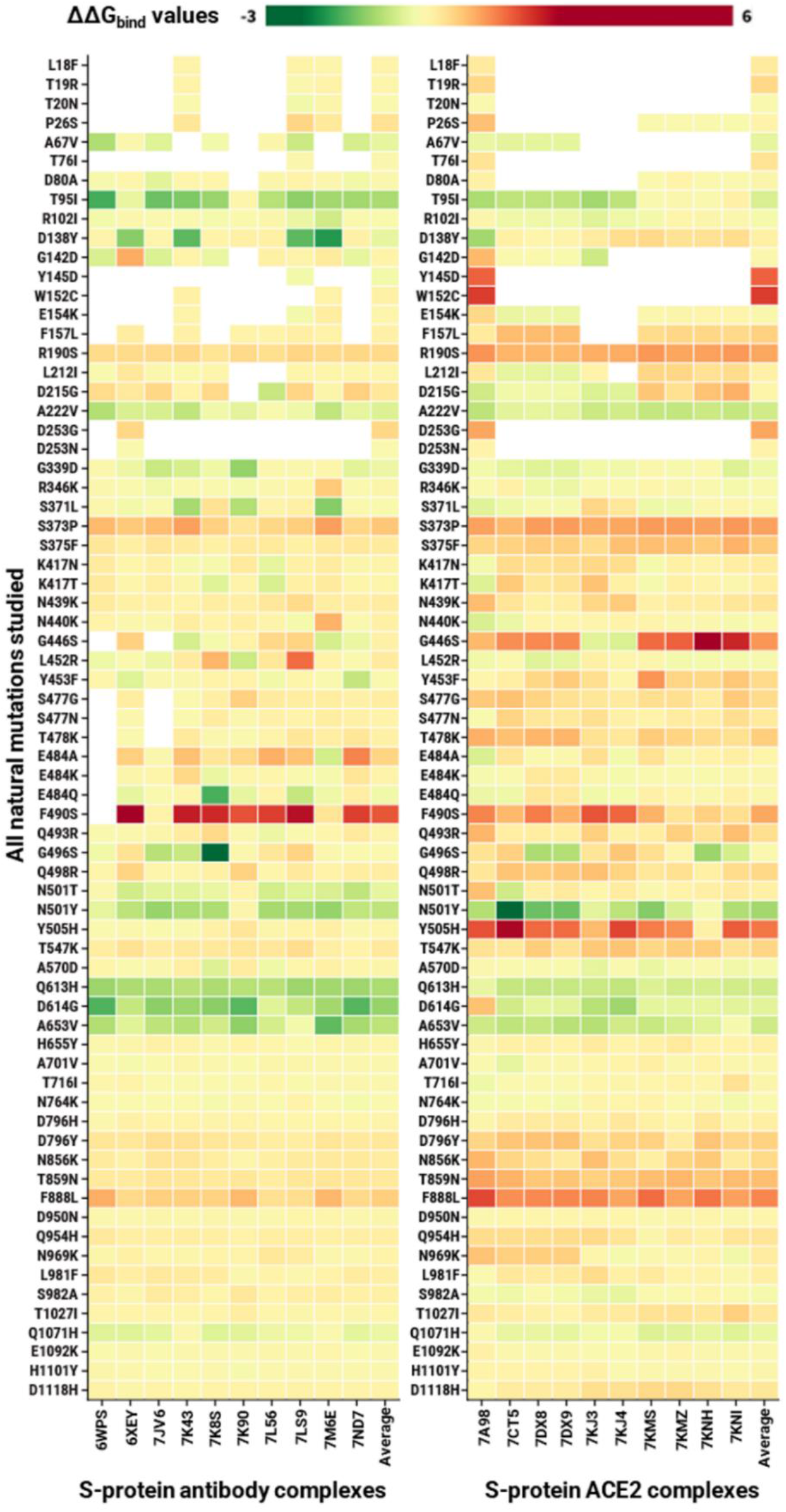
Heatmap of changes in binding affinity estimated for the studied natural mutations. The mutations with comparatively increased binding affinity are shown in green, whereas mutations that reduce binding affinity relative to the Wuhan reference strain are shown in red. Mutations in sites that were not available in the experimental structures are shown in white.

Interestingly, D614G showed increased affinity towards both antibodies and ACE2, but more towards the antibodies. This would be consistent with a relative loss of fitness in a population where these antibodies circulated, i.e., the antibodies were made to bind the S-protein with this very early fixated mutation present. The literature suggests that the D614G variant increases infectivity but reduces the affinity for ACE2 because of a faster dissociation rate.^70^ Increased selectivity for the antibody ensemble was observed for T95I, D138Y, R190S, S373P, S375F, K417N, K417T, N439K, G446S, Y453F, S477G, S477N, T478K, E484Q, Q493R, Q498R, Y505H, T547K, Q613H, D614G, A653V, D796H, D796Y, N856K, T859N, F888L, Q954H, N969K, L981F, T1027I, and D1118H. Approximately half of these are located on the RBD. These observations are consistent with the studied set of human monoclonal antibodies having been developed to target the circulating natural mutations.

In contrast, mutations such as D215G, A222V, E484A, F490S, N501Y, A570D and S982A showed increased selectivity for ACE2. Comparatively, a very small number of mutations exhibit increased affinity for ACE2, and these might exhibit the antibody escape tendency. The N501Y mutant is reported to display 5-10 times higher binding to ACE2 than wild-type.^57, 71–73^ These mutations might be suspected to induce a future selective advantage by having more antigenic drift vs. the studied antibodies.

### ΔΔG_bind_ variations due to S-protein structure and types of mutations

As mentioned in the Methods section, the mutation dataset was divided into nine groups (**Table S4**). **Figures S4-S5** show the detailed comparison of these groups for each of the 20 used experimental structures. In general, natural mutations exhibited significantly less reduced affinity compared to the control group of random mutations and even all possible hypothetical mutations located at the binding interface in both AB and ACE2 sets. When averaging the 10 structures and using group comparison (Figure 3 and **Figure S6)**, a similar trend was observed for the nine mutation groups. However, the effect of interface and delta mutations was highly variable in the AB dataset. Interface residues showed the lowest average ΔΔG_bind_ compared to any other mutation group. Of the nine groups, delta mutations exhibited the highest average affinity change, followed by gamma, whereas omicron showed similarly decreased affinity in both datasets. For the antibody escape mutations, we observed a similar lowered binding, consistent with substantial antibody escape.

**Figure 3.**
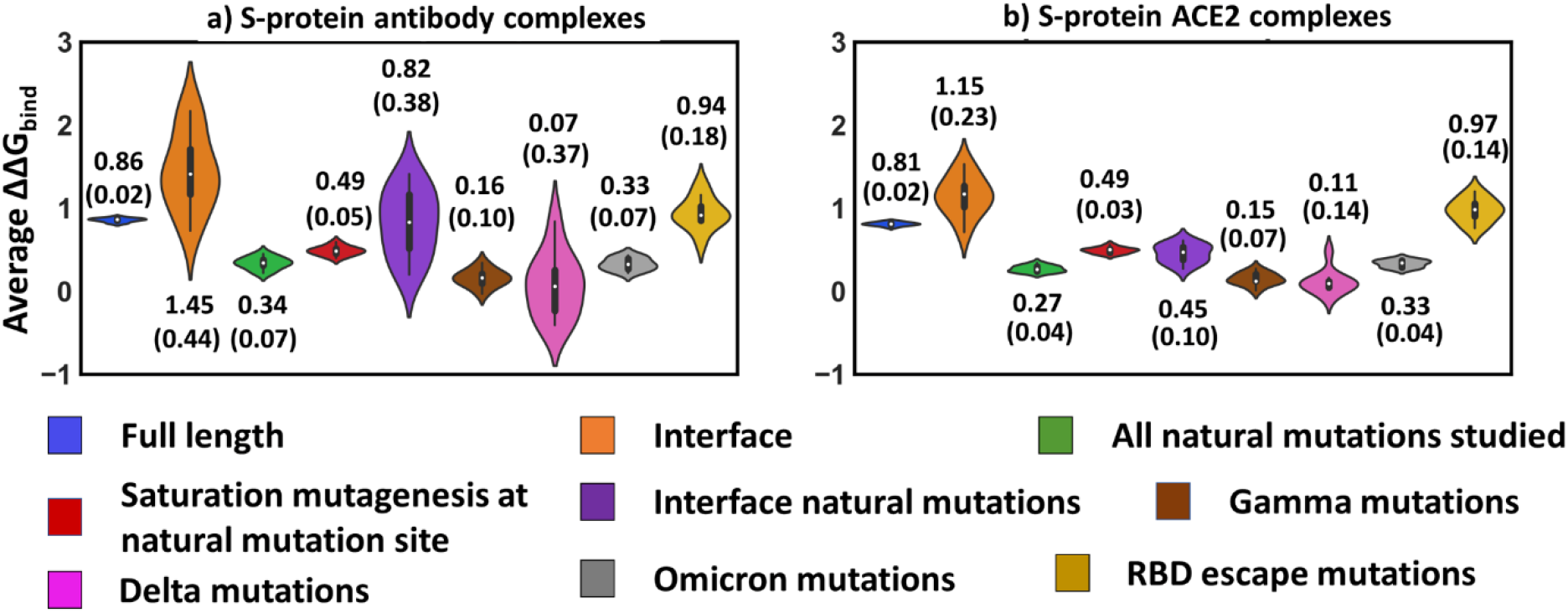
Average change in S-protein’s binding affinity (ΔΔG_bind_) to antibodies and ACE2 for the studied mutation categories, averaged over the 10 structures. The average and standard deviation values (in brackets) are shown for each mutation group. A positive value indicates a decreased binding and *vice-versa*.

Figure 3 overall indicates that natural mutations had higher affinity impacts compared to random mutations in the full S-protein, including interface sites separately. Also, the natural mutations in the binding interface showed decreased affinity compared to full group of natural mutations, including gamma, delta, and omicron groups. We note that the standard deviations (SD) for the mutation groups are higher for the AB dataset than for ACE2, consistent with the larger expected structural epitope variance of the antibodies, whereas ACE2 represents one type of protein receptor only. This heterogeneity also explains why results using just one antibody as representative of antigenic drift is not meaningful.

Figure 4 reveals differences between the behavior of the mutation groups for the two datasets: (i) random mutations in full length S-protein show significantly different binding affinity changes (blue), i.e., lower binding for AB structures; (ii) the interface region exhibits comparatively more variable and low average affinity in AB structures (orange), though the difference is not significant; (iii) the natural mutations show significantly lower antibody binding (green), which may relate to their AB escape tendency; (iv) similar prominent effects were observed for the interface natural mutations, i.e. lower AB binding. All these observations suggest weaker binding of the S-protein to antibody structures for natural and random mutations. However, no significant differences were observed for the individual comparisons between gamma, delta, omicron, and escape mutations. This might be due to the smaller dataset for comparison because a significant difference was noted for the combined natural mutation datasets.

**Figure 4.**
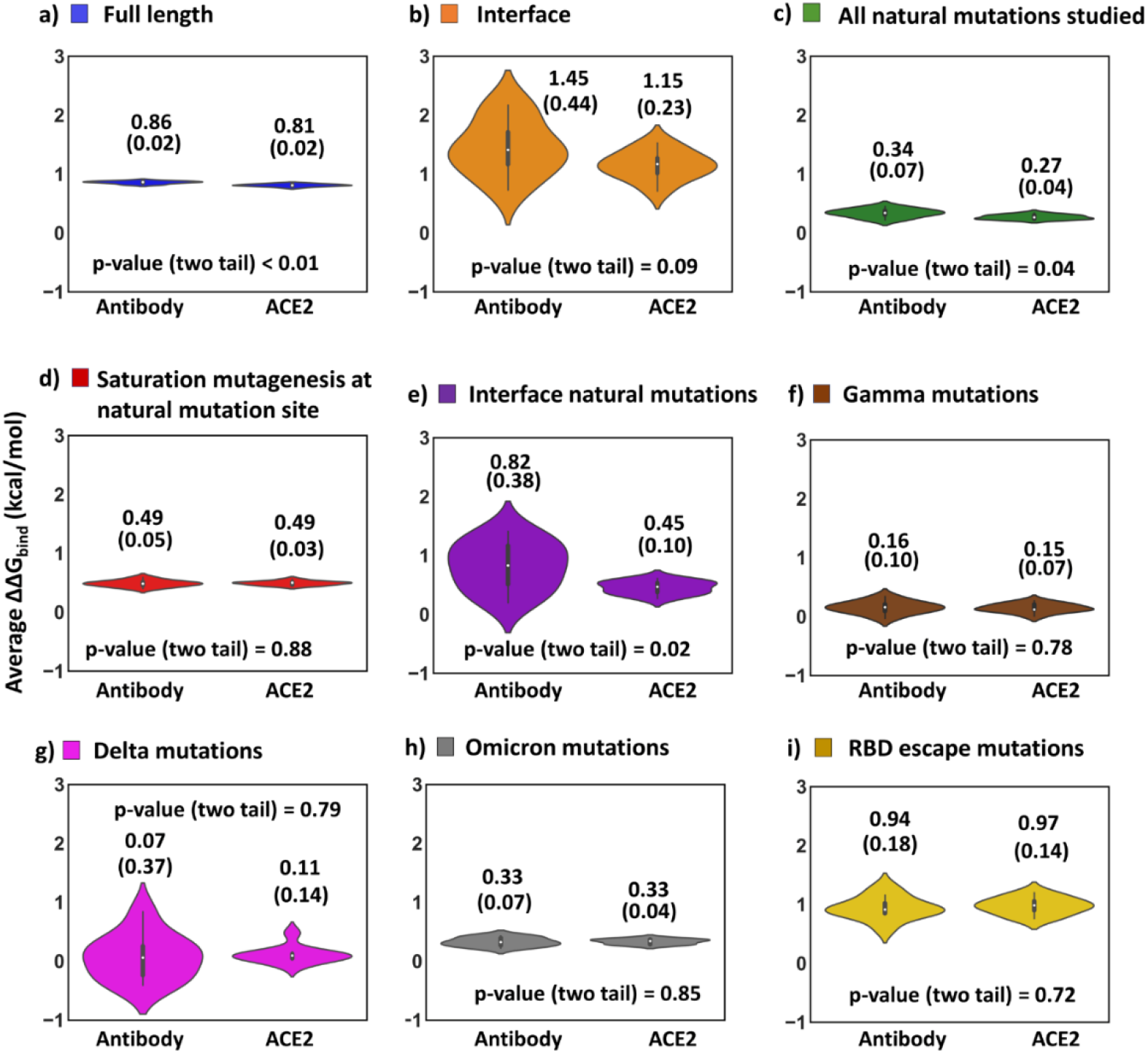
Comparison of binding affinity changes for S-protein binding to antibody or ACE2. Average ΔΔG_bind_ value, standard deviation (in bracket), and p-value (two tail) in each subplot. A p-value smaller than 0.05 was used to indicate high probability (95% confidence level) that the mean mutation effects differ for the two types of protein targets.

Overall, the interface natural mutations had the largest significant change in selectivity (binding energy difference between the two protein targets), 0.82 vs. 0.45 kcal/mol, as indicated in Figure 4e, where these positive values indicate reduced binding to both with respect to wild-type, but more reduced binding to AB than ACE2. This indicates that the interface natural mutations have comparatively more fitness towards ACE2. The group of all possible and all natural mutations also showed a similar significant effect of comparatively reduced antibody binding suggesting viral fitness for ACE2.

We further studied the conformational dependence of the binding affinity changes by dividing structures into subsets based on the number of AB/ACE2 binding partners and RBD conformation as detailed in **Supplementary section 3**. A very similar effect of each of these structural subsets was found, which is consistent with the combined effect of the ten AB/ACE2 structures, indicating that the results are not influenced by the change in the number of binding partners or different RBD conformations.

### Estimating fitness effects by a simple model

The calculated change in fitness values (ΔF, equation 3) were plotted in a heatmap (Figure 5 and **S9**) for all natural mutations studied for each ACE2 and antibody-bound complex. ΔF was calculated in three ways: Figure 5a displays ΔF as the difference between average ΔΔG_bind_ for 10 structures in ACE2 set and ΔΔG_bind_ for each structure in the AB set; similarly, Figure 5b shows ΔF as the difference between ΔΔG_bind_ for each structure in ACE2 set and the average ΔΔG_bind_ for 10 structures in AB set (X-axis shows the PDB id; the average column in the average value for the 10 structures); whereas Figure 5c shows average ΔF as the difference in the average ΔΔG_bind_ values of 10 structures between each ACE2 and AB sets.

**Figure 5.**
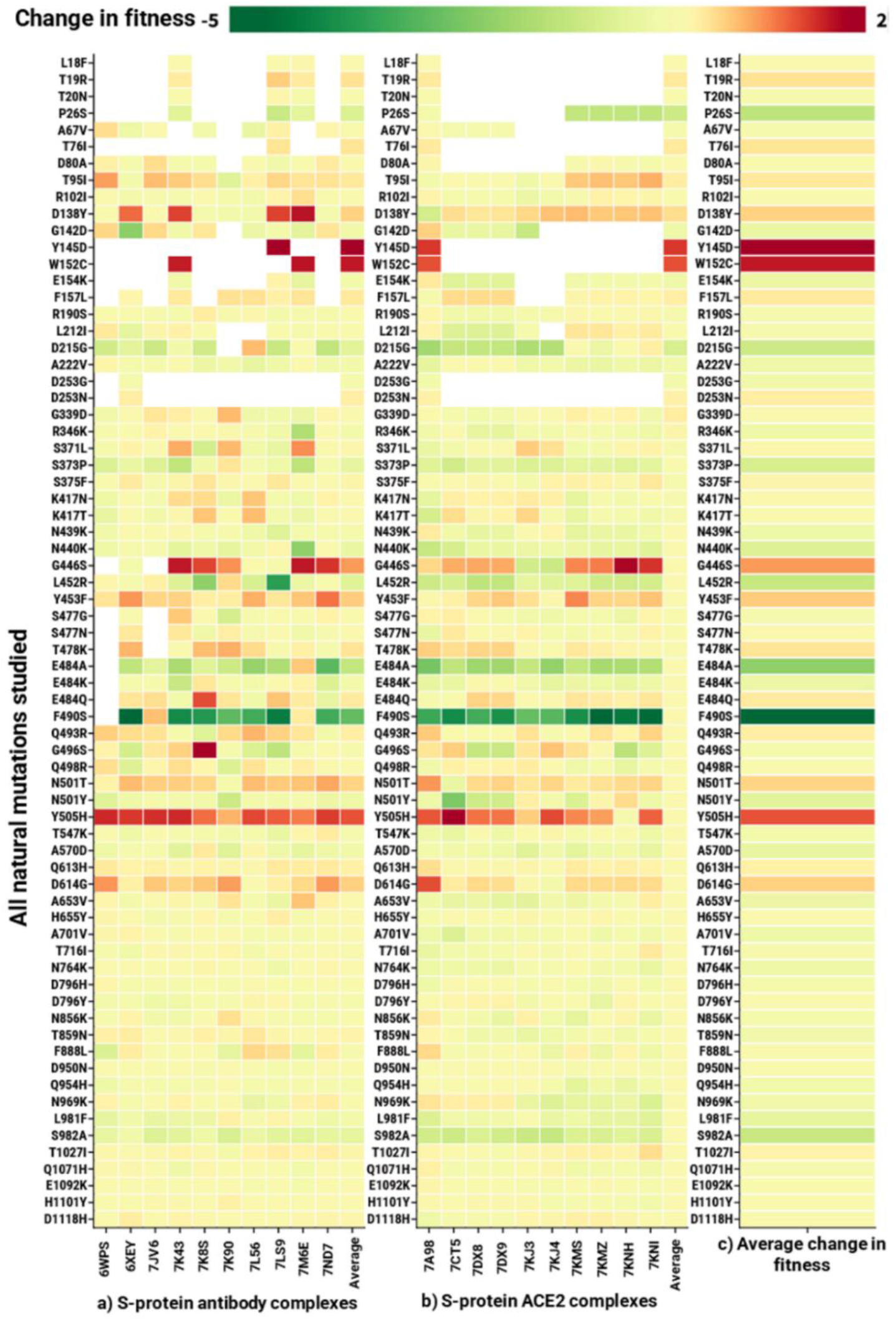
Heatmap showing the estimated change in fitness using a simple two-property fitness model. **(a)** ΔF = Average ΔΔG_bind(MUT, 10 ACE2)_ – ΔΔG_bind(MUT, each AB)_. The x-axis shows the PDB ID of the antibody set, with values being averages of 10 structures. **(b)** ΔF = ΔΔG_bind(MUT, each ACE2)_ – Average ΔΔG_bind(MUT, 10 AB)_. The x-axis shows the PDB ID of the ACE2 set, with values being averages of 10 structures. **(c)** Average ΔF = Average ΔΔG_bind(MUT, 10 ACE2)_ – Average ΔΔG_bind(MUT, 10 AB)_.

Clearly, these individual mutations and average values in each Figure 5a**, 5b** and **5c** differ by the choice of structural set used for computation. Therefore, relying on one structure is not sufficient for drawing any conclusions. The consensus obtained in Figure 5c is of interest because it considers variable antibody epitope and structural conformations. Here, a more negative (green) value indicates more ACE2 binding or more antibody escape, whereas a more positive (red) value shows that the mutation increases binding to antibodies relative to ACE2. Residues P26S, G142D, E154K, D215G, N440K, S373P, L452R, E484A, E484K, F490S, N501Y, A570D, L981F, and S982A have more affinity for ACE2 receptor (cutoff < −0.25). These variations might lead to increased viral fitness and cause antibody escape. T19R, T76I, T95I, D138Y, Y145D, W152C, G446S, Y453F, T478K, N501T, Y505H, and D614G reveal a less binding affinity for ACE2 but more binding for antibodies (cutoff > 0.25). This mutation set reveals higher antibody binding that may neutralize the viral particles.

This simple fitness estimate, the difference between change in binding of ACE2 and antibody, can be used to anticipate important mutations leading to increased or decreased viral fitness. This initial computational effort to model fitness is further validated by comparison with experimental data below.

### Validation of the approach

To validate our computational strategy, we compared our results with the three experimental datasets (0-2) defined in **Table S11**. The experimental ACE2 binding data^57^ (**dataset 0**) was compared with the computed ΔΔG_bind_ for ACE2 (**Figure S10**). The average ΔΔG_bind_ for mutations in ACE2 complexes showed a highly significant correlation with R = 0.37 (p < 0.01), and when using binned method with 0.25 binned data, the correlation increased to R = 0.84 (p < 0.01) (**Figure S10**), as reported earlier.^58^ This indicating the experimental ACE2 binding data are in parts defined by our computations. It is important to note here that three experimental datasets are highly squeezed with very less difference between several data points. However, comparing while taking average of small intervals in the data defined as bins may be more suitable for any comparison, as reported earlier.^59^ Therefore, we used the binned strategy (0.25 bins) for all further comparisons.

**Figure S11** shows the comparison of the experimental ACE2 binding (**dataset 0**) versus −log (antibody escape fractions) (**datasets 1** and **2**). Clearly, the direct correlations (R = 0.84 for dataset 0 vs. 1, and R = 0.56 for dataset 0 vs. 2 using 0.25 bins) confirm our argument about the generic effects of mutations in protein-protein interactions, i.e., mutation that tend to decrease S-protein affinity for one type of protein (ACE2) also reduces binding for the other type of protein (say antibodies).

The second approach involved comparison of two sets of experimental AB escape fraction data (**datasets 1** and **2**; **Table S11**) with our computed AB binding effects. Each of the datasets 1 and 2 was comprised of the average effect of ten antibodies. The average ΔΔG_bind_ for each mutation was correlated with experimental values for AB average mutation escape.^74, 75^ Computed change in binding correlates significantly with mutation escape data with the same and meaningful direction (**Figure S12a;** dataset 1, R = 0.25 and p <0.01; dataset 2, R = 0.21 and p <0.01). Interestingly, the correlation increased to R = 0.85 (p <0.01) and 0.60 (p = 0.06) for datasets 1 and 2 using 0.25 bins (**Figure S12a**). We also specifically checked the correlation of the AB-bound S-proteins present in our dataset with the experimental AB escape data (**Figure S12b**). We observed a good correlation for experimental mutation escape of C002 antibody with calculated ΔΔG_bind_ for the C002 PDB structure (PDB code 7K8S; R= 0.66, p-value = 0.02 for 0.25 bins) with similar direction of regression line as with datasets 1 and 2. Similar trend was observed for C144 antibody correlation (PDB code 7K90; R = 0.65, p-value = 0.01 for 0.25 bins) (**Figure S12b**). This suggests that our computations partially explain the mutation escape data.

Most importantly, in the third approach, the experimental differences (ACE2 – AB), using **Table S11** data, were compared with computed change in fitness for the RBD mutations (Figures 6 and **S13**). Interestingly, for both comparisons (ACE2 – AB datasets 1 and 2), experimental differences correlated significantly with computed differences, which was a fair comparison because the computed fitness is the difference between change in S-protein binding to ACE2 and AB (i.e. ACE2 – AB). The experimental values are the difference between ACE2 binding and −log (AB escape fraction). The binned data showed good correlations with R values of 0.68 (p-value <0.01) and 0.87 (p-value <0.01) for datasets 1 and 2 respectively (Figure 6). The directions of correlations were meaningful and in accordance with the experimental datasets, as the computed fitness value decreased (more ACE2 binding), the experimental difference increased (more ACE2 binding), thus validating our approach.

**Figure 6.**
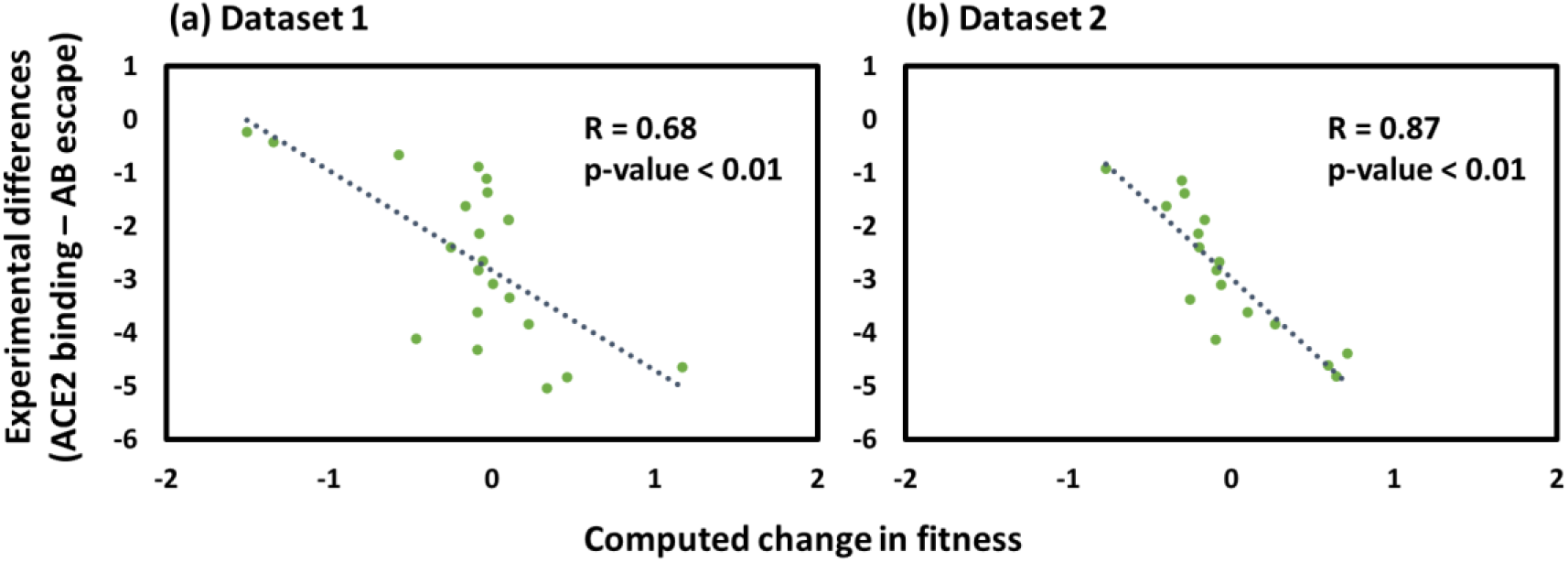
Scatter plots for the comparison of computed change in fitness with experimental differences (ACE2 binding – AB escape) Experimental data were binned into 0.25 bins and correlated with the computed data. **(a)** Dataset 1. **(b)** Dataset 2.

Overall, our computed binding data was able to partially reproduce the effect shown by diverse experimental studies on ACE2 and antibodies (**Figures S10-S13**), indicating the potential of our approach in computing binding effects, and therefore the fitness.

### Considering intraprotein epistasis

In order to understand the effect (including non-additive effects, i.e., “intraprotein epistasis”) of multiple mutations simultaneously present in the S-protein, we compared ΔΔG (kcal/mol) calculated using FoldX^60^ for different variants with all their mutations present simultaneously, as seen in the variants (Figure 7a). We observed that for gamma variants (G1 and G2), omicron and RBD escape show resemblance in stability, with apo S-protein being the most stable and ACE2 complexes being the least stable. However, for the delta variant, the ACE2 complexes were more stabilized than the AB complexes with the least stability change for apo S-protein. In the same analysis, we also compared the stability of different variants for apo and holo structures (**Figure S14**). We found that the delta variant has high stability in AB- and ACE2-bound complexes, whereas RBD escape mutation decreases the stability of the protein in apo and holo forms. In addition, omicron mutations are decreasing stability of ACE2 complexes largely compared to AB complexes.

**Figure 7.**
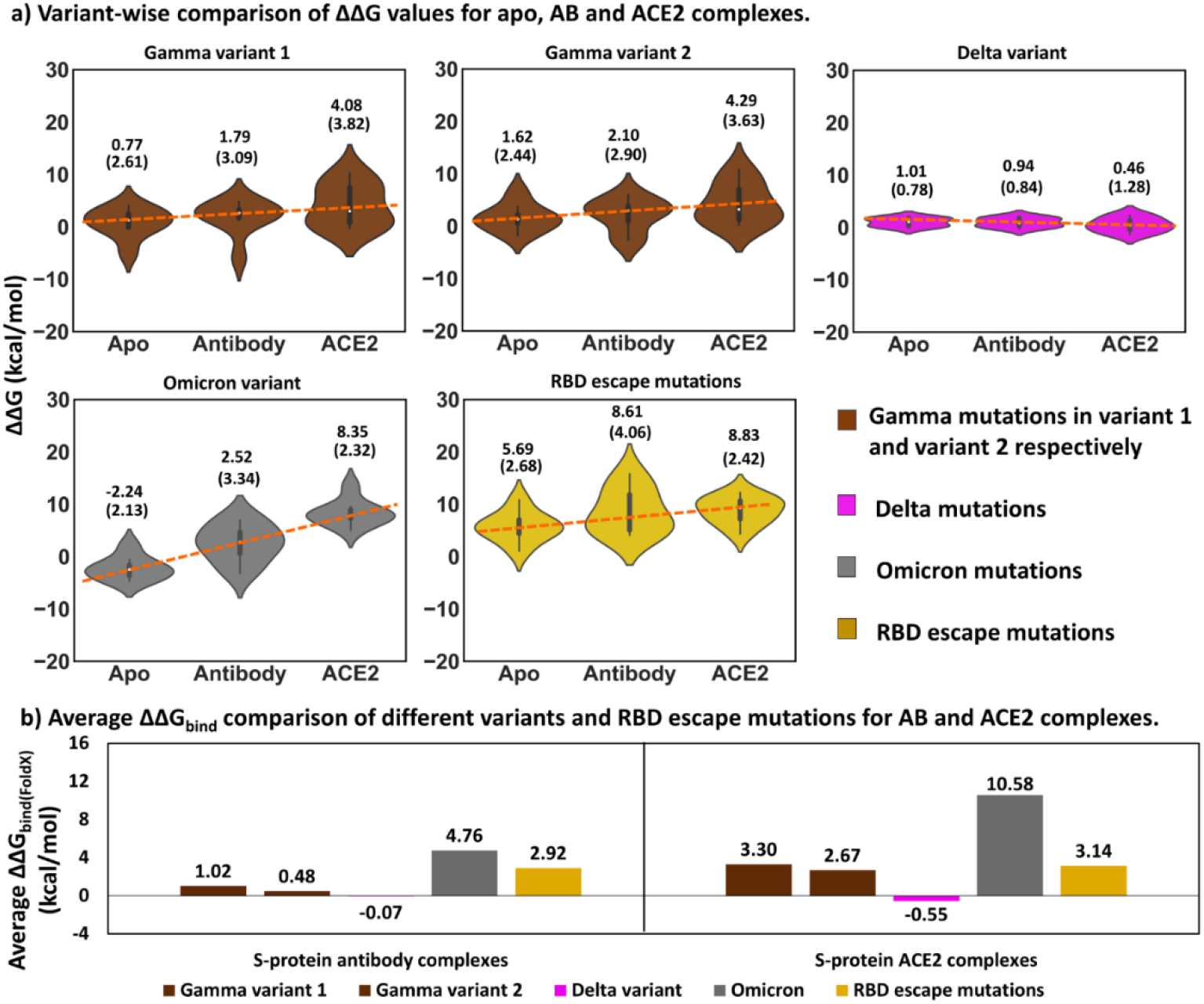
Summarizing epistasis in relation to ACE2 and AB binding. **(a)** ΔΔG values of apo, AB- and ACE2-bound complexes were compared for different variants using epistasis effect. **(b)** Comparing average change in binding free energy of different variants upon epistasis for AB-bound and ACE2-bound complexes. Positive ΔΔG show destabilizing and negative stabilizing effect of mutation. Similarly, positive ΔΔG_bind_ represents decreased binding.

When comparing the change in binding free energy (ΔΔG_bind(FoldX)_ = average ΔΔG_(AB/ACE2)_ – average ΔΔG_(apo)_) value for AB vs. ACE2 caused by mutation, a similar pattern was seen for the variants and RBD escape mutations (Figure 7b). A negative ΔΔG_bind(FoldX)_ value indicates increased binding affinity and *vice-versa*. The average values suggest that delta mutations stabilize both AB and ACE2 complexes, whereas omicron mutations tended to do the opposite, but the net effect is again the most important from a fitness perspective, as discussed above.

### Limitations and strengths of the method

The fitness function defined here is a proof of the concept and not exact. It shows the ability of our approach to model fitness exceptionally well, realising the fact that some variables contributing to fitness were kept constant; still the simple two-property model can reproduce the effect, which is the novelty of our work. The model did not include the roles of innate and adaptive immunity in infection clearance, the evolved viral mechanisms for reduced detection,^76, 77^ and the dynamics of an immune response.^78^ The advanced model might be expected to include such immune factors. However, our simplified model could partly reproduce the effects of several experimental datasets. It is an initial step towards using protein structure-based tools for making fitness functions at the virus organism level.

The model is based on the ten antibodies dataset binding to the RBD region similar to the ten ACE2 dataset binding. Such approach ensured consistency to reduce noise and maintain uniformity in computations. We do not deny the possibility of including more antibodies binding to multiple regions of S-protein, which is not considered here, since this may lead to heterogeneity of the approach. Some antibodies may not compete with ACE2 for binding to the S-protein yet still neutralize the virus.^41, 79, 80^ Most of the experimental AB escape fraction data used for validating our approach are different from the set of computed AB data used in the present study, still the comparisons showed good correlations with consistent and meaningful direction of regression line, indicating the strength of our approach.

The experimental binding assay data and the experimental structures do not represent accurately the in vivo S-protein structure, which is heavily glycosylated.^81^ Even beyond this glycan effect, the cryo-EM structures may not reflect the physiologically most representative conformation state due to the exposure of the samples to the cryo-temperature.^23, 24, 82^ The rapid cooling may freeze some conformational dynamics,^83–91^ and some of the conformations in the S-protein may be pH- or temperature dependent in ways that will add complexity to real-world applications of the results.^92^ Also, the experimental assays additionally are done on single mutants and thus do not reflect the same effects in variants where some introduced mutation may correlate (same-gene epistasis).

The use of differences in binding affinity caused by mutation, and differences of these between ACE2 and antibodies reduce systematic errors considerably, and the use of structure-averaged mean effects reduce structural noise and increase precision, which partly explain the robustness of our results. However, we note that systematic effects due to glycosylation could change conclusions reported both here and in the experimental assays, so the work here should be seen as a starting point for establishing fitness estimates using molecular modeling, not as final best estimates.

## Conclusions

We have described a simple model and a computational protocol to estimate the fitness of arising mutations in the SARS-CoV-2 virus, based on the structure-based computational estimates of S-protein binding to a cocktail of important antibodies and human ACE2, emphasizing that fitness relates to a competitive binding situation, where the virion seeks to bind ACE2 before it is bound to circulating antibodies. The model uses many protein structures to compute robust estimates of these effects and takes advantage of a large degree of systematic error cancellation by considering differences between differences in estimated binding affinities (changes in ACE2–AB binding selectivity). Thus, while the model and protocol still have some limitations, it represents the best model so far in our estimate on a path towards predicting and rationalizing the impact of arising mutations in SARS-CoV-2 at the molecular level, work that is still in progress and requires much expansion before being applicable to e.g., other viruses as well.

## Supplementary information

Details about the work are provided in the supplementary file. It contains details about the antibody and ACE2 datasets used in the study, group of mutations, conformational dependence of the binding affinity changes, experimental ACE2/antibody binding/escape datasets for validation, detailed statistical tests (violin plots, correlation plots and t-test results), and epistatic analyses.

## Acknowledgments

RM acknowledges SERB-SRG, Govt of India for funding via grant SRG/2022/000304-C. RM, ST acknowledge IIT Bhilai for supporting this work via Research Initiation Grant (RIG), number 2005900. ST acknowledges the Ministry of Education, Government of India, for PMRF fellowship grant number 5001800.

## Author contributions

RM and KPK conceptualized the work. RM supervised the work. ST performed the work. ST, RM and KPK wrote and refined the manuscript.

