## Supplementary file for "Predicting Virus Fitness: Towards a structure-based computational model"

*Rukmankesh Mehra: 0000-0001-6010-1514*

*Kasper P. Kepp: 0000-0002-6754-7348*

*Shivani Thakur: 0000-0002-7869-6648*

**1. Details about S-protein-antibodies structural datasets**

The AB complexes (**Figure S1**) have the RBD in the down-conformation and comprise overlapping S-protein binding sites with ACE2. Residue-specific details regarding the ABs and S-protein binding regions are shown in **Tables S5-S6** and discussed below:

S309 (present in 6WPS) potently neutralizes authentic SARS-CoV2 virus ($IC_{50}$ = 79 ng/mL) via recognizing a conserved N343 glycan on the RBD of S-protein[1]. S309 binds to both open and close state of glycoprotein without competing with the ACE2 receptor[1].

The fragment antibody (Fab) of human AB 2-4 (present in 6XEY) binds the RBD and competes with ACE2 for binding[2]. It locks the RBD in the down conformation and neutralizes SARS-CoV2 with an estimated $IC_{50}$ value of 57 ng/mL[2,3].

S2H13 (present in 7JV6) is known to neutralize SARS-CoV2 S-protein ($IC_{50}$ = 500 ng/mL) by recognizing an RBD epitope accessible in both open and closed state of S-protein and blocking ACE2 attachement[4]. Both ACE2 and S2H13 bind to overlapping epitopes[4].

C002 (present in 7K8S) and C144 (present in 7K90) recognizes both up and down conformations of the RBD and various residues close to the RBD[5]. C002 shows neutralizing activity against SARS-CoV-2 pseudovirus with $IC_{50}$ = 8 ng/mL[5,6] whereas C144 shows neutralization against authentic SARS-CoV-2 with an $IC_{50}$ value of 2.55 ng/mL[6].

COVOX-316 (present in 7ND7) binds to all down conformation of S-protein targeting RBM and blocks ACE2 attachment[7]. It neutralizes SARS-CoV2 S-protein with $IC_{50}$ = 10 ng/mL[8].

AB 1-57 (present in 7LS9) binds to RBD by competing with the ACE2 receptor and neutralizes live virus with $IC_{50}$ = 8 ng/mL[9]. It shows small quaternary interaction with neighboring RBDs via recognizing N165 glycan[9].

S2M11 (present in 7K43) neutralizes authentic SARS-COV2 with $IC_{50}$ = 3 ng/mL by competitively blocking ACE2 attachment and locking the S-protein in a closed state via recognizing quaternary epitope through electrostatic interactions[10].

BG 10-19 (present in 7M6E) locks the S-protein trimer in a distinct closed state that cannot bind ACE2 and potently neutralizes SARS-CoV2 with $IC_{50}$ = 9 ng/mL[11]. Its neutralization mechanism involves recognition of N165 glycan on adjacent N-terminal domain accessible in both up/down conformation and formation of quaternary epitope interactions between two adjacent RBDs restricting "up" conformation of RBD[11].

Fab 2-43 (present in 7L56) binds to the RBM in the down state of S-protein only and recognizes N343 glycans on neighboring RBDs, forming strong quaternary interactions to lock the RBD in the down conformation, thereby inhibiting ACE2 interaction[12]. It neutralizes live SARS-CoV2 virus with an $IC_{50}$ value of 3 ng/mL[2].

Overall, the studied structures contain distinct antibodies with varying accessibility to the RBD epitopes, various VH and VL segment genes, and CDRH3 and CDRL3 length, and also diverse neutralizing activity. There are similarities between some of the ABs, such as (a) S309, S2M11, C144, Fab 2-43 recognize N343 glycan for binding, which may lock RBD into closed conformation, blocking its attachment to the ACE2 receptor, (b) S2M11, C144, COVOX-316, Fab 2-4, Fab 2-43 and 1-57 blocks all RBD in down conformation and binds to RBM overlapping region[12,8], (c) VH 1-2 ABs (Fab 2-4, S2M11, Fab 2-43) have similar binding conformations to an epitope overlapping with the ACE2 binding site[12]. C144 shows a similar angle approach, and epitope to VH 1-2 ABs[12], (d) S309, S2H13, C002, C144 and BG 10-19 recognize RBD in both up and down conformation, (e) S2M11, C144, BG 10-19 and Fab 2-43 have similar interactions with two neighboring RBDs and form quaternary interactions, (f) S309, Fab 2-4, 1-57, S2H13, COVOX-316 target only one RBD and do not interact with neighboring RBDs, and (g) C002, as a special case, shows quaternary interaction with a downwards oriented RBD only if an adjacent RBD is at the same time in an up-conformation.

### 2. Details about groups of mutations

The group of all natural mutations (**Table S4**) comprises the data of gamma, delta, and omicron mutations, i.e., set $c$ = sets $f + g + h$. The interface natural mutations ($e$) is a subset of ($b$) and ($c$). The mutation data were compiled from UniProt[13], CDC[14], and the WHO[15] websites (accessed on April 13, 2022).

We included the AB escape data to analyze the effect of the mutations in AB binding. The escape mutation data was taken from two datasets (**Table S7**). From Baum et al.[16], log (average $IC_{50}$) values were used as a criteria to distinguish strong escape mutations. In case of AB binding to measure SARS-CoV2 neutralization, a larger $IC_{50}$ value indicates less AB binding (more antibody escape). Similarly, larger average $IC_{50}$ also indicates more AB escape. Therefore, a larger log($IC_{50}$) of the mutation will correspond to a higher escape tendency of the mutation from that particular antibody. We thus converted $IC_{50}$ values for WT and mutations into the log(average $IC_{50}$) and found that for wild-type (WT), log (average $IC_{50}$) is −10.2. For strong escape mutations, log(average $IC_{50}$) [mutant >> WT], i.e., AB shows less binding and more escape compared to WT. Based on these calculations, we selected seven mutations having log (average $IC_{50}$) in the range of −9 to −10 (>> −10.21), considered to be strong RBD escape mutations.

From the data by Zost et al.[17], we used the average response of five antibodies to select prominent escape mutations, normalized to WT response = 1, for 15 mutations showing reduced monoclonal AB binding. In this study, high response corresponds to more AB binding for that mutation. We calculated the average response using the response value of five antibodies, and mutations with an average response of less than 1 were selected as RBD escape mutations (total 6 mutations) i.e., average response [WT >> mutant] indicates less AB binding and more escape to mutant residue as compared to WT. Based on these data, a total of 13 mutations were selected as RBD escape mutations and analyzed separately as a validation test set.

#### 3. Conformational dependence of the binding affinity changes

To better understand the conformational dependence of the $\Delta\Delta G_{bind}$, we divided the S-protein-ACE2 structures into 3 ACE2 (5 structures) and 2 ACE2 (5 structures) bound, and all RBD up (6 structures) and 2 RBD up (4 structures) conformations. The average $\Delta\Delta G_{bind}$ values for each of the nine groups were computed for these structural subsets, in addition to the combined effect of 10 structures (**Figure S7**). We found a very similar effect of each of these structural subsets, which is consistent with the combined effect of 10 structures, indicating that the results are not influenced by the change in the number of binding partners or different RBD conformations.

Similarly, the S-protein AB structures were also classified based on the number of AB units interacting with each RBD, by dicing into structures having one AB unit (6 structures) or two AB units (4 structures) (**Figure S8**), which also indicated that this had a minor influence on the results relative to the larger and more direct impact of the mutation itself, probably because any specific mutating site rarely interacts with the same site in the other unit in the structures with multiple antibodies present. Structural comparison of the 10 S-protein structures bound to ACE2 showed RMSD of 1.6−3.2 Å with regards to the highest-resolution structure 7KJ4 (3.4 Å resolution) (**Table S13**). However, for the antibody dataset, all S-protein structures were in RBD down conformations. The RMSD of these structures was in the range 2.3–4.2 Å with regards to the structure 7K43 (2.60 Å resolution) (**Table S13**). The RMSD difference between two high-resolution S-protein structures (from this study), one from each AB- and ACE2-bound group (7K43 vs. 7KJ4) was 1.99 Å.

**Table S1. S-protein-AB complexes used in the present study (AB dataset). The RBD segments of three S-protein chains are in down conformations in all structures.**

| PDB | Resolution (Å) | % Outliers | AB present in the structure | Number of AB units | Reference |
|---|---|---|---|---|---|
| 6WPS | 3.1 | 0.3 | S309 | 3[a] | Pinto et al.[1] |
| 6XEY | 3.3 | 0.1 | Fab 2-4 | 3[a] | Liu et al.[2] |
| 7JV6 | 3.0 | 0.3 | S2H13 | 3[a] | Piccoli et al[4] |
| 7K8S | 3.4 | 0.2 | C002 | 3[a] | Barnes et al.[5] |
| 7ND7 | 3.6 | 0.4 | COVOX-316 | 3[a] | Dejnirattisai et al.[7] |
| 7LS9 | 3.4 | 0.1 | 1-57 | 3[a] | Cerutti et al.[9] |
| 7K43 | 2.6 | 0.2 | S2M11 | 3[b] | Tortorici et al.[10] |
| 7K90 | 3.2 | 0.1 | C144 | 3[b] | Barnes et al.[5] |
| 7M6E | 3.3 | 0.2 | BG 10-19 | 3[b] | Scheid et al.[11] |
| 7L56 | 3.6 | 0.2 | Fab 2-43 | 3[b] | Rapp et al.[12] |

[a] - Complexes in which 1 AB unit (one heavy & one light chain) interacts with each S-protein RBD region.

[b] - Complexes where 2 AB units interact with each S-protein RBD region.

**Table S2. S-protein-ACE2 complexes used in the present study (ACE2 dataset). The ACE2 units are bound on RBD up conformations.**

| PDB | Resolution (Å) | % Outliers | RBD conformation | Number of ACE2 bound | Reference |
|---|---|---|---|---|---|
| 7A98 | 5.4 | 0.2 | 3 up | 3 | Benton et al.[18] |
| 7CT5 | 4.0 | 0.1 | 3 up | 3 | Guo et al.[19] |
| 7KJ4 | 3.4 | 0.4 | 3 up | 3 | Xiao et al.[20] |
| 7KMS | 3.6 | 0.0 | 3 up | 3 | Zhou et al.[21] |
| 7KNI | 3.9 | 0.0 | 3 up | 3 | Zhou et al.[21] |
| 7DX9 | 3.6 | 0.3 | 3 up | 2 | Yan et al.[22] |
| 7KJ3 | 3.7 | 0.4 | 2 up, 1 down | 2 | Xiao et al.[20] |
| 7DX8 | 2.9 | 0.3 | 2 up, 1 down | 2 | Yan et al.[22] |
| 7KMZ | 3.6 | 0.0 | 2 up, 1 down | 2 | Zhou et al.[21] |
| 7KNH | 3.7 | 0.0 | 2 up, 1 down | 2 | Zhou et al.[21] |

**Table S3. Overview of used structures, showing the number of residues in each chain of the S-protein and the antibodies.**

| PDB ID of AB-bound complexes | Number of residues in each S-protein chain | Number of residues in each AB chain | | PDB ID of ACE2-bound complexes | Number of residues in each S-protein chain |
|---|---|---|---|---|---|
| | | Heavy | Light | | |
| **6WPS** | 955, 955, 955 | 102, 102, 102 | 123, 123, 123 | **7A98** | 1066, 1066, 1066 |
| **7K8S** | 1004, 1004, 1004 | 125, 125, 125 | 107, 107, 107 | **7KMS** | 1033, 1033, 1033 |
| **7K90** | 1000, 1000, 1000 | 129, 129, 129 | 108, 108, 108 | **7KNI** | 1030, 1030, 1030 |
| **7LS9** | 1097, 1097, 1097 | 129, 129, 129 | 108, 108, 108 | **7DX9** | 1006, 1006, 1007 |
| **7M6E** | 1034, 1034, 1034 | 123, 123, 123 | 108, 108, 108 | **7CT5** | 1006, 1007, 1006 |
| **6XEY** | 1034, 1034, 1030 | 122, 122, 122 | 108, 108, 108 | **7KMZ** | 1007, 1030, 1031 |
| **7JV6** | 977, 977, 977 | 118, 118, 118 | 109, 109, 109 | **7KNH** | 1007, 1030, 1031 |
| **7L56** | 983, 992, 994 | 129, 129, 129 | 110, 110, 110 | **7DX8** | 1007, 971, 1006 |
| **7ND7** | 1002, 1002, 1002 | 122, 122, 122 | 109, 109, 109 | **7KJ3** | 981, 981, 961 |
| **7K43** | 1034, 1034, 1034 | 122, 122, 122 | 105, 105, 105 | **7KJ4** | 981, 981, 981 |

**Table S4. Mutation groups (or sets) defined in this work.**

| Group | Mutations belonging to each group |
|---|---|
| ***(a)*** **Full length** | All possible mutations in S-protein ACE2 or S-protein AB complex. |
| ***(b)*** **Interface residues** | S-protein ACE2 complex: R403, D405, K417, V445, G446, Y449, Y453, L455, F456, Y473, A475, G476, S477, N481, E484, G485, F486, N487, Y489, F490, Q493, S494, G496, Q498, T500, N501, G502, V503, Y505 |
| | S-protein AB complex: N334, L335, P337, G339, E340, F342, N343, A344, T345, R346, Y351, N354, K356, R357, S359, N360, S366, V367, L368, N370, S371, A372, S373, F374, K417, W436, N440, L441, K444, V445, G446, G447, N448, Y449, N450, L452, Y453, L455, F456, T470, E471, N481, G482, V483, E484, G485, F486, N487, Y489, F490, L492, Q493, S494, G496, Q498, P499, T500, R509 |
| ***(c)*** **All natural mutations studied** | L5F, S13I, L18F, T19R, T20N, P26S, A67V, G75V, T76I, D80A, T95I, R102I, D138Y, G142D, Y145D, W152C, E154K, F157L, R190S, L212I, D215G, A222V, D253G/N, G339D, R346K, S371L, S373P, S375F, K417N/T, N439K, N440K, G446S, L452R, Y453F, S477G/N, T478K, E484K/A/Q, F490S, Q493R, G496S, Q498R, N501Y/T, Y505H, T547K, A570D, Q613H, D614G, A653V, H655Y, Q677H, N679K, P681H/R, A701V, T716I, N764K, D796H/Y, N856K, T859N, F888L, D950N, Q954H, N969K, L981F, S982A, T1027I, Q1071H, E1092K, H1101Y, D1118H, V1176F, G1219V |
| ***(d)*** **Saturation mutagenesis at natural mutation site** | 19 other possible mutations at the natural mutation site. |
| ***(e)*** **Interface natural mutations** | S-protein ACE2 complex: K417N/T, G446S, Y453F, S477G/N, E484K/A/Q, F490S, Q493R, G496S, Q498R, N501Y/T, Y505H |
| | S-protein AB complex: G339D, R346K, S371L, S373P, K417N/T, N440K, G446S, L452R, Y453F, E484K/A/Q, F490S, Q493R, G496S, Q498R |
| ***(f)*** **Gamma (P.1)** | L18F, T20N, P26S, D138Y, R190S, K417N, K417T, E484K, N501Y, D614G, H655Y, T1027I, V1176F |
| ***(g)*** **Delta (B.1.617.2)** | T19R, L452R, T478K, D614G, P681R, D950N |
| ***(h)*** **Omicron (B.1.1.529)** | A67V, T95I, Y145D, L212I, G339D, S371L, S373P, S375F, K417N, N440K, G446S, S477N, T478K, E484A, Q493R, G496S, Q498R, N501Y, Y505H, T547K, D614G, H655Y, N679K, P681H, N764K, D796Y, N856K, Q954H, N969K, L981F |
| ***(i)*** **RBD escape mutations** | Q321L, V341I, N354D, Q409E, A435S, I472V, Y508H, K444A, G447R, F486A, N487A, P499R, N501A |

**Table S5. Residues assigned to the interface in the used structures.**

| S.no. | PDB IDs of S-protein AB complexes | Interface residues present in the respective PDB structure | PDB IDs of S-protein ACE2 complexes | Interface residues present in the respective PDB structure |
|---|---|---|---|---|
| 1. | **6WPS** | N334, L335, P337, G339, E340, N343, A344, T345, R346, N354, K356, R357, S359, N360, N440, L441, K444 | **7A98** | D405, Y449, Y453, L455, F456, Y473, A475, G476, S477, G485, F486, N487, Y489, F490, Q493, G496, Q498, T500, N501, G502, V503, Y505 |
| 2. | **6XEY** | G446, Y449, Y453, L455, F456, V483, E484, G485, F486, Y489, F490, Q493, S494, G496, Q498 | **7CT5** | R403, K417, Y449, Y453, L455, F456, A475, G476, S477, F486, N487, Y489, Q493, S494, G496, Q498, T500, N501, G502, V503, Y505 |
| 3. | **7JV6** | G447, Y449, N481, G482, V483, E484, G485, F486, F490, S494, Q498 | **7DX8** | R403, K417, Y449, F456, A475, G476, S477, F486, N487, Y489, F490, Q493, G496, Q498, T500, N501, G502, V503, Y505 |
| 4. | **7K8S** | A372, S373, N440, K444, V445, G447, Y449, N450, L452, Y453, L455, T470, E471, N481, G482, V483, E484, G485, F486, Y489, F490, L492, Q493, S494, G496, T500 | **7DX9** | R403, K417, Y449, F456, A475, G476, S477, N481, F486, N487, Y489, Q493, G496, Q498, T500, N501, G502, V503, Y505 |
| 5. | **7K43** | G339, F342, N343, T345, V367, L368, S371, A372, S373, F374, W436, N440, L441, K444, G446, Y449, L452, L455, F456, E484, G485, F486, Y489, F490, Q493, S494, G496, Q498 | **7KJ3** | R403, K417, V445, Y449, L455, F456, Y473, A475, G476, E484, G485, F486, N487, Y489, F490, Q493, G496, Q498, T500, N501, G502, V503, Y505 |
| 6. | **7K90** | G339, F342, N343, V367, L368, N370, S371, A372, S373, F374, K417, W436, N440, Y449, N450, L455, F456, V483, E484, G485, F486, N487, Y489, F490, Q493, S494, Q498 | **7KJ4** | R403, K417, V445, G446, Y449, L455, F456, A475, G476, G485, F486, N487, Y489, F490, Q493, S494, G496, Q498, T500, N501, G502, V503, Y505 |
| 7. | **7L56** | S366, N370, G446, G447, N448, Y449, N450, L455, F456, V483, E484, G485, F486, Y489, F490, L492, Q493, S494, Q498 | **7KMS** | R403, K417, G446, Y449, Y453, L455, F456, Y473, A475, G476, S477, E484, F486, Y489, Q493, G496, Q498, T500, N501, G502, V503, Y505 |
| 8. | **7LS9** | R346, Y351, K444, V445, G446, G447, N448, Y449, N450, L452, T470, E484, G485, F490, L492, Q493, S494, Q498 | **7KMZ** | R403, K417, G446, Y449, Y453, L455, F456, Y473, A475, G476, F486, Y489, Q493, G496, Q498, T500, N501, G502, V503, Y505 |

| | | | | |
|---|---|---|---|---|
| 9. | **7M6E** | G339, F342, N343, T345, R346, V367, L368, S371, S373, F374, W436, N440, L441, K444, V445, G446, N448, Y449, N450, E484, G485, F486, N487, G496, Q498, P499, T500, R509 | **7KNH** | R403, K417, G446, Y449, L455, F456, Y473, A475, G476, G485, F486, Y489, Q493, G496, Q498, T500, N501, G502, V503, Y505 |
| 10. | **7ND7** | G446, Y449, Y453, L455, F456, V483, E484, G485, F486, N487, Y489, F490, Q493, S494, G496, Q498 | **7KNI** | R403, K417, G446, Y449, Y453, L455, F456, Y473, A475, G476, S477, F486, Y489, F490, Q493, G496, Q498, T500, N501, G502, V503, Y505 |

**Table S6. Common interface residues of the S-protein in the studied ten AB- and ten ACE2-bound complexes.**

| |
|---|
| K417, V445, G446, Y449, Y453, L455, F456, N481, E484, G485, F486, N487, Y489, F490, Q493, S494, G496, Q498, T500 |

**Table S7. Experimental AB data used to define RBD escape mutations.**

| Dataset name | For each mutation | | | Data value for wild-type | Range of data used in selection | No. of mutations selected |
|---|---|---|---|---|---|---|
| | Data available | Data conversion | Data significance | | | |
| **Baum et al.**[16] | $IC_{50}$ for 8 ABs | Log (average $IC_{50}$) | $\text{Log(average IC}_{50}) \propto \frac{\text{AB escape}}{\text{AB binding}}$ | Log (average $IC_{50}$) = -10.21 | Data value between -9 and -10 | 7 |
| **Zost et al.**[17] | Response (normalized to wild-type) for 5 ABs | Average response | $\text{Average response} \propto \frac{\text{AB binding}}{\text{AB escape}}$ | Average response = 1 | Average response < 1 | 6 |

**Table S8. P-values obtained from t-tests (testing for same structure-averaged mean $\Delta\Delta G_{bind}$) for various combinations of AB- versus ACE2-bound S-protein structures.**

| Variable 1 (AB-bound complexes) | Variable 2 (ACE2-bound complexes) | p-value (One-tail) | p-value (Two-tail) |
|---|---|---|---|
| **1. Full length** | **Full length** | 0.0007 | 0.001 |
| **2. Interface** | **Interface** | 0.04 | 0.09 |
| **3. All natural mutations studied** | **All natural mutations studied** | 0.02 | 0.04 |
| **4. Saturation mutagenesis at natural mutation site** | **Saturation mutagenesis at natural mutation site** | 0.44 | 0.88 |
| **5. Interface natural mutations** | **Interface natural mutations** | 0.01 | 0.02 |
| **6. Gamma** | **Gamma** | 0.39 | 0.78 |
| **7. Delta** | **Delta** | 0.39 | 0.79 |
| **8. Omicron** | **Omicron** | 0.43 | 0.85 |
| **9. RBD escape mutations** | **RBD escape mutations** | 0.36 | 0.72 |

**Table S9. P-values obtained from t-tests of antibody complexes (testing for same structure-averaged mean $\Delta\Delta G_{bind}$) to determine whether the mutation groups overall behave differently.**

| Variable 1 | Variable 2 | p-value (one tail) | p-value (two tail) |
|---|---|---|---|
| 1. **Full length** | Interface | 0.002 | 0.003 |
| 2. **Full length** | All natural mutations studied | 1.81E-09 | 3.62E-09 |
| 3. **Full length** | Saturation mutagenesis at natural mutation site | 4.12E-09 | 8.23E-09 |
| 4. **Full length** | Interface natural mutations | 0.39 | 0.77 |
| 5. **Full length** | Gamma | 1.07E-09 | 2.13E-09 |
| 6. **Full length** | Delta | 6.02E-05 | 0.00012 |
| 7. **Full length** | Omicron | 2.68E-09 | 5.35E-09 |
| 8. **Full length** | RBD escape mutations | 0.11 | 0.22 |
| 9. **Interface** | All natural mutations studied | 9.84E-06 | 1.97E-05 |
| 10. **Interface** | Saturation mutagenesis at natural mutation site | 3.46E-05 | 6.93E-05 |
| 11. **Interface** | Interface natural mutations | 0.0001 | 0.0003 |
| 12. **Interface** | Gamma | 7.84E-06 | 1.57E-05 |
| 13. **Interface** | Delta | 2.06E-05 | 4.13E-05 |
| 14. **Interface** | Omicron | 7.28E-06 | 1.46E-05 |
| 15. **Interface** | RBD escape mutations | 0.0003 | 0.0006 |
| 16. **All natural mutations studied** | Saturation mutagenesis at natural mutation site | 1.07E-07 | 2.14E-07 |
| 17. **All natural mutations studied** | Interface natural mutations | 0.0008 | 0.002 |
| 18. **All natural mutations studied** | Gamma | 0.001 | 0.002 |
| 19. **All natural mutations studied** | Delta | 0.02 | 0.04 |
| 20. **All natural mutations studied** | Omicron | 0.20 | 0.40 |
| 21. **All natural mutations studied** | RBD escape mutations | 6.71E-07 | 1.34E-06 |
| 22. **Saturation mutagenesis at natural mutation site** | Interface natural mutations | 0.007986 | 0.015971 |
| 23. **Saturation mutagenesis at natural mutation site** | Gamma | 9.19E-06 | 1.84E-05 |
| 24. **Saturation mutagenesis at natural mutation site** | Delta | 0.002344 | 0.004687 |
| 25. **Saturation mutagenesis at natural mutation site** | Omicron | 9.09E-07 | 1.82E-06 |
| 26. **Saturation mutagenesis at natural mutation site** | RBD escape mutations | 8.04E-06 | 1.61E-05 |
| 27. **Interface natural mutations** | Gamma | 0.0006 | 0.001 |
| 28. **Interface natural mutations** | Delta | 1.7E-05 | 3.39E-05 |
| 29. **Interface natural mutations** | Omicron | 0.00076 | 0.001 |
| 30. **Interface natural mutations** | RBD escape mutations | 0.12 | 0.24 |
| 31. **Gamma** | Delta | 0.28 | 0.57 |
| 32. **Gamma** | Omicron | 0.002 | 0.004 |
| 33. **Gamma** | RBD escape mutations | 1.95E-06 | 3.9E-06 |
| 34. **Delta** | Omicron | 0.02 | 0.05 |
| 35. **Delta** | RBD escape mutations | 2.69E-05 | 5.38E-05 |
| 36. **Omicron** | RBD escape mutations | 3.96E-07 | 7.93E-07 |

**Table S10. P-values obtained from t-tests of ACE2 complexes (testing for same structure-averaged mean $\Delta\Delta G_{bind}$) to determine whether the mutation groups overall behave differently.**

| Variable 1 | Variable 2 | p-value (one tail) | p-value (two tail) |
|---|---|---|---|
| 1. **Full length** | Interface | 0.0009 | 0.002 |
| 2. **Full length** | All natural mutations studied | 1.41E-10 | 2.82E-10 |
| 3. **Full length** | Saturation mutagenesis at natural mutation site | 4.04E-09 | 8.08E-09 |
| 4. **Full length** | Interface natural mutations | 2.38E-06 | 4.75E-06 |
| 5. **Full length** | Gamma | 3.07E-10 | 6.15E-10 |
| 6. **Full length** | Delta | 1.34E-07 | 2.68E-07 |
| 7. **Full length** | Omicron | 5.23E-10 | 1.05E-09 |
| 8. **Full length** | RBD escape mutations | 0.003 | 0.005 |
| 9. **Interface** | All natural mutations studied | 2.46E-07 | 4.92E-07 |
| 10. **Interface** | Saturation mutagenesis at natural mutation site | 2.24E-06 | 4.47E-06 |
| 11. **Interface** | Interface natural mutations | 1.42E-07 | 2.84E-07 |
| 12. **Interface** | Gamma | 1.75E-07 | 3.49E-07 |
| 13. **Interface** | Delta | 1.02E-07 | 2.05E-07 |
| 14. **Interface** | Omicron | 4.57E-07 | 9.14E-07 |
| 15. **Interface** | RBD escape mutations | 0.01 | 0.03 |
| 16. **All natural mutations studied** | Saturation mutagenesis at natural mutation site | 1.81E-10 | 3.61E-10 |
| 17. **All natural mutations studied** | Interface natural mutations | 8.59E-05 | 0.0002 |
| 18. **All natural mutations studied** | Gamma | 0.0003 | 0.0006 |
| 19. **All natural mutations studied** | Delta | 0.002 | 0.003 |
| 20. **All natural mutations studied** | Omicron | 6.64E-06 | 1.33E-05 |
| 21. **All natural mutations studied** | RBD escape mutations | 3.5E-08 | 7E-08 |
| 22. **Saturation mutagenesis at natural mutation site** | Interface natural mutations | 0.099 | 0.199 |
| 23. **Saturation mutagenesis at natural mutation site** | Gamma | 4.08E-08 | 8.15E-08 |
| 24. **Saturation mutagenesis at natural mutation site** | Delta | 5.23E-06 | 1.05E-05 |
| 25. **Saturation mutagenesis at natural mutation site** | Omicron | 6.55E-08 | 1.31E-07 |
| 26. **Saturation mutagenesis at natural mutation site** | RBD escape mutations | 5.96E-07 | 1.19E-06 |
| 27. **Interface natural mutations** | Gamma | 7.21E-06 | 1.44E-05 |
| 28. **Interface natural mutations** | Delta | 5.85E-05 | 0.0001 |
| 29. **Interface natural mutations** | Omicron | 0.001 | 0.003 |
| 30. **Interface natural mutations** | RBD escape mutations | 1.19E-06 | 2.37E-06 |
| 31. **Gamma** | Delta | 0.28 | 0.57 |
| 32. **Gamma** | Omicron | 7.02E-06 | 1.4E-05 |
| 33. **Gamma** | RBD escape mutations | 5.94E-09 | 1.19E-08 |
| 34. **Delta** | Omicron | 0.0003 | 0.0007 |
| 35. **Delta** | RBD escape mutations | 4.9E-08 | 9.8E-08 |
| 36. **Omicron** | RBD escape mutations | 1.5E-07 | 3E-07 |

**Table S11. Description of the experimental dataset used for calculating correlation with our computations. The common data points between experimental and computed data were used for correlation analysis.**

| Dataset | AB/ ACE2 data | Number of AB | Data type available for RBD mutation | Data values after conversion (used for each mutation) | Data points for comparison | Data significance | Reference |
|---|---|---|---|---|---|---|---|
| **0** | ACE2 | - | Binding average | Binding average | 3283 | Binding average ∝ ACE2 binding | Starr et al.[23] |
| **1** | AB | 10 | Escape fraction | $-\log_{10}$(average escape fraction of 10 ABs) | 1924 | $-\log_{10}$(average escape fraction) ∝ AB binding | Greaney et al.[24] |
| **2** | AB | 10 | Escape fraction | $-\log_{10}$(average escape fraction of 10 ABs) | 2346 | $-\log_{10}$(average escape fraction) ∝ AB binding | Greaney et al.[25] |

**Table S12. Mutation categories to study the effect by introducing all the mutations of the group together using FoldX (instead of point mutation as studied above).**

| | |
|---|---|
| **Gamma variant 1** | L18F, T20N, P26S, D138Y, R190S, K417N, E484K, N501Y, D614G, H655Y, T1027I, V1176F |
| **Gamma variant 2** | L18F, T20N, P26S, D138Y, R190S, K417T, E484K, N501Y, D614G, H655Y, T1027I, V1176F |
| **Delta variant** | T19R, L452R, T478K, D614G, P681R, D950N |
| **Omicron** | A67V, T95I, Y145D, L212I, G339D, S371L, S373P, S375F, K417N, N440K, G446S, S477N, T478K, E484A, Q493R, G496S, Q498R, N501Y, Y505H, T547K, D614G, H655Y, N679K, P681H, N764K, D796Y, N856K, Q954H, N969K, L981F |
| **RBD escape mutations** | Q321L, V341I, N354D, Q409E, A435S, I472V, Y508H, K444A, G447R, F486A, N487A, P499R, N501A |

**Table S13. RMSD values calculated with regards to 7K43 and 7KJ4 (the structures having the highest resolution) for AB and ACE2-bound complexes, respectively. RMSD value between the two highest resolution structures 7K43 (AB) vs. 7KJ4 (ACE2) was 1.99 Å.**

| PDB ID of AB-bound complexes | RMSD value (w.r.t. 7K43 in Å) | PDB ID of ACE2-bound complexes | RMSD value (w.r.t. 7KJ4 in Å) |
|---|---|---|---|
| **6WPS** | 2.694 | **7A98** | 2.673 |
| **7K8S** | 2.332 | **7KMS** | 2.343 |
| **7K90** | 3.152 | **7KNI** | 2.711 |
| **7LS9** | 2.695 | **7DX9** | 2.765 |
| **7M6E** | 2.510 | **7CT5** | 2.998 |
| **6XEY** | 3.007 | **7KMZ** | 2.678 |
| **7JV6** | 2.459 | **7KNH** | 2.954 |
| **7L56** | 4.172 | **7DX8** | 3.208 |
| **7ND7** | 3.210 | **7KJ3** | 1.617 |

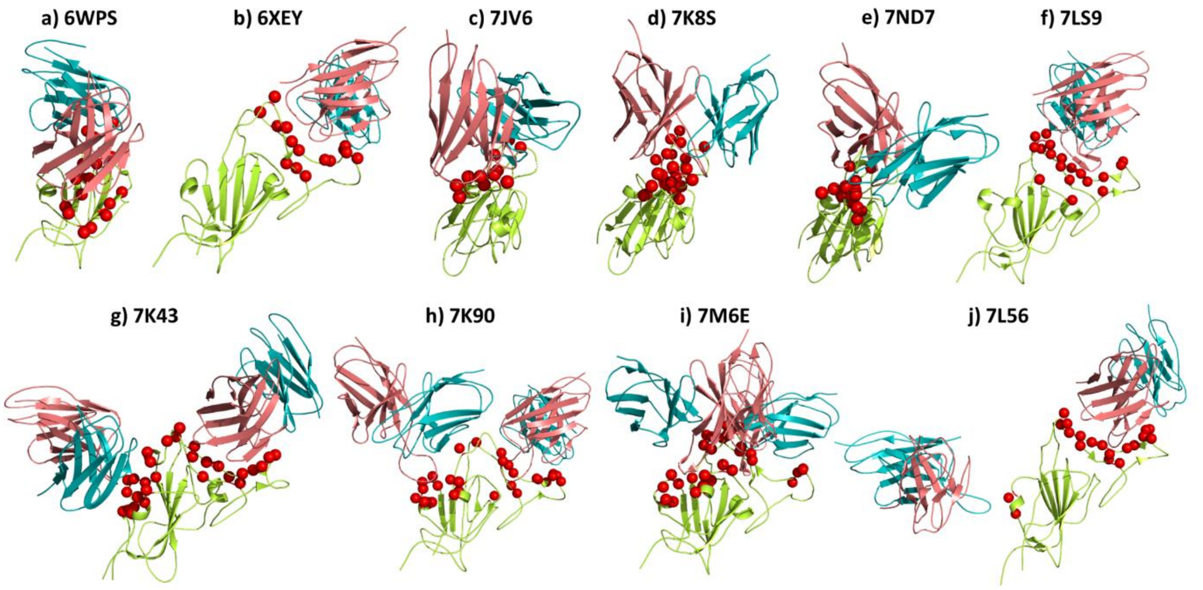


**Figure S1. S-protein-AB complexes studied in this work, with the interface residues represented as red balls.** **(a)** 6WPS by Pinto et al.[1], **(b)** 6XEY by Liu et al.[2], **(c)** 7JV6 by Piccoli et al.[4], **(d)** 7K8S by Barnes et al.[5], **(e)** 7ND7 by Dejnirattisai et al.[7], **(f)** 7LS9 by Cerutti et al.[9], **(g)** 7K43 by Tortorici et al.[10], **(h)** 7K90 by Barnes et al.[5], **(i)** 7M6E by Scheid et al.[11] and **(j)** 7L56 by Rapp et al.[12] .Only the RBD region of S-protein is shown (green color); the S-protein-AB interacting residues are shown in red, the heavy chain of AB in pink, and the light chain in blue.

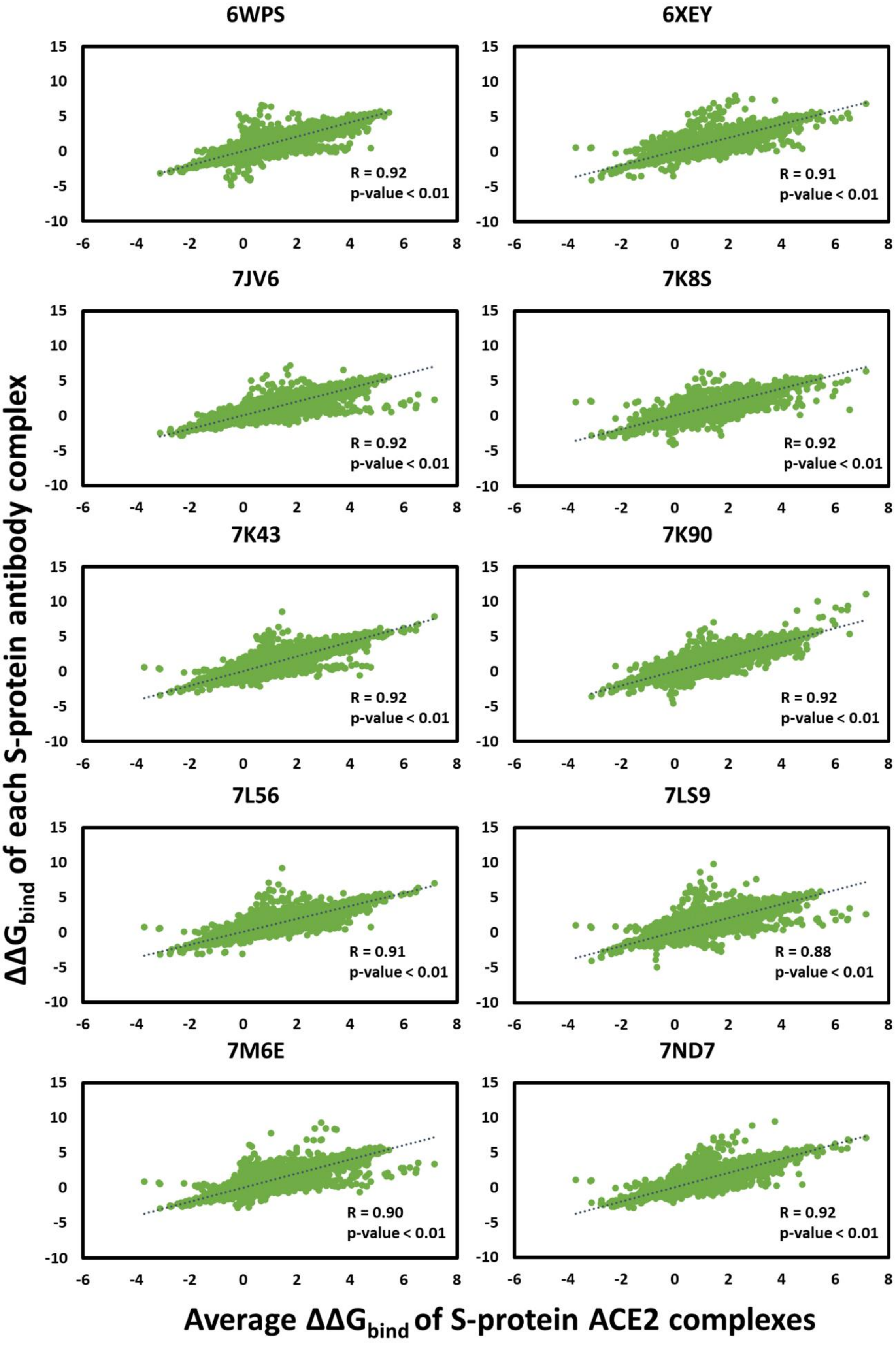


**Figure S2. Scatter plot for comparison of $\Delta\Delta G_{bind(each\ AB)}$ with average $\Delta\Delta G_{bind(10\ ACE2)}$ complex.** The Pearson correlation coefficients (R) and p-values are shown for each plot.

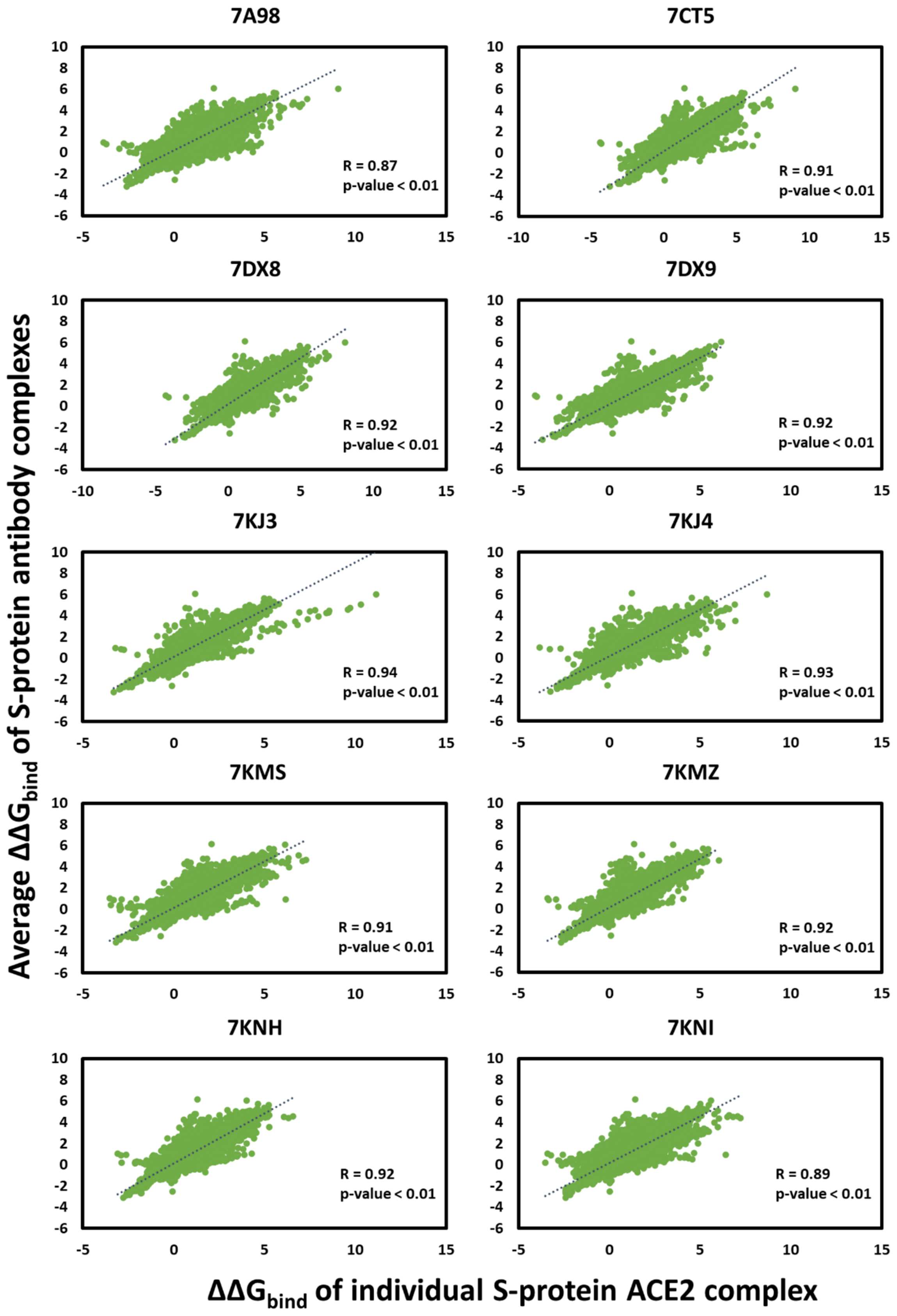


**Figure S3. Scatter plot for comparison of $\Delta\Delta G_{bind(each\ ACE2)}$ with average $\Delta\Delta G_{bind(ten\ AB)}$ complex.** The Pearson correlation coefficients (R) and p-values are shown for each plot.

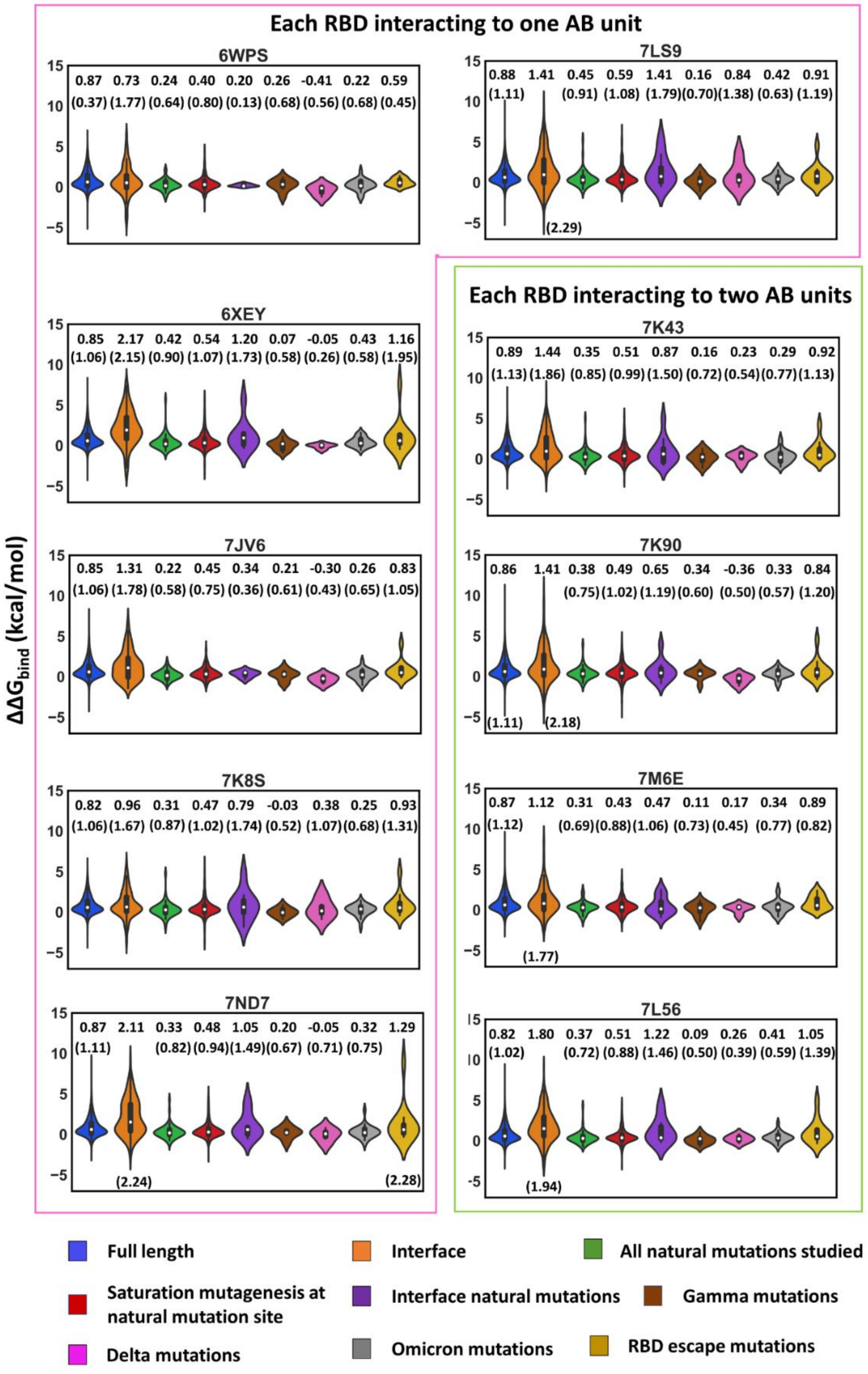


**Figure S4. Change in binding affinity for AB-bound complexes compared for different mutation groups.** The pink boxes represent $\Delta\Delta G_{bind}$ values of complexes with each RBD of S-protein interacting to one AB unit (one heavy and one light chain). The green boxes represent $\Delta\Delta G_{bind}$ values of complexes in which each RBD is interacting with two AB units. Average $\Delta\Delta G_{bind}$ was calculated for each mutation group. Standard deviations are shown in brackets. Violin plots were generated using Python libraries Matplotlib, Seaborn, and Pandas.

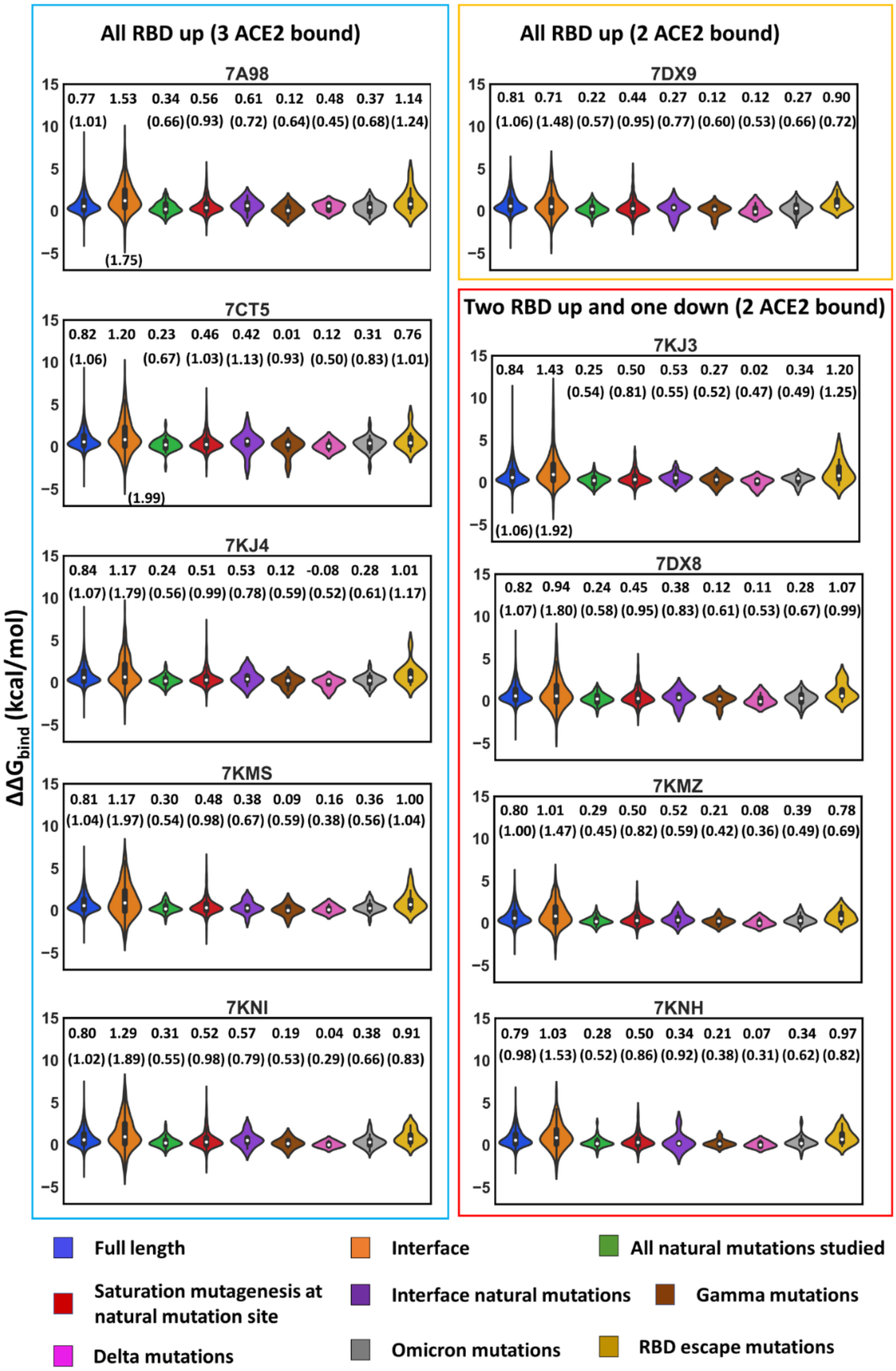


**Figure S5. Binding affinities of S-protein to ACE2 for each mutation category, highlighting mutations in concerning variants.** $\Delta\Delta G_{bind}$ plots in the blue box represent five complexes with all RBD in up-conformation and 3 ACE2 units bound to it. The orange box represents the $\Delta\Delta G_{bind}$ for all S-protein RBDs in up-conformation but only two ACE2 units bound. The red box represents $\Delta\Delta G_{bind}$ of four complexes with two RBDs in up-conformation, one in down conformation, and two ACE2 bound to each up-RBD. The figure shows the average $\Delta\Delta G_{bind}$ values for each mutation category and standard deviations in brackets.

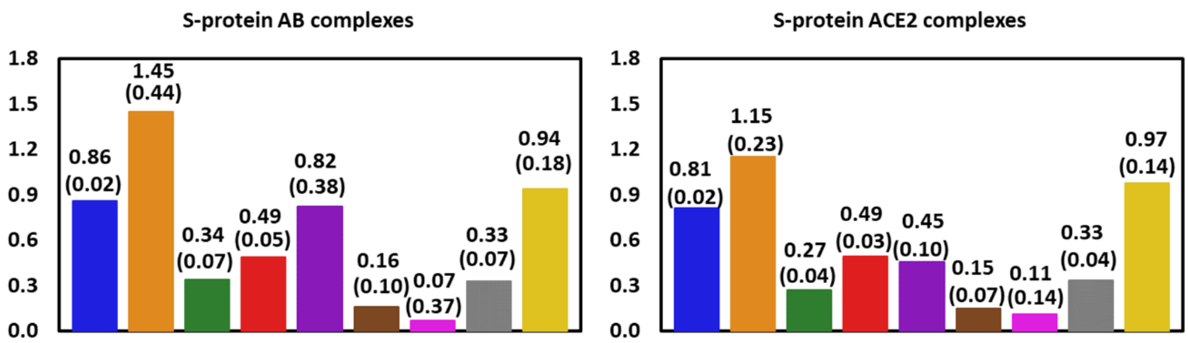


**Figure S6. Comparison of the average change in S-protein's binding affinity ($\Delta\Delta G_{bind}$) of ABs and ACE2 for the studied mutation categories, averaged over used structures.** Standard deviation values are written in brackets for each mutation group.

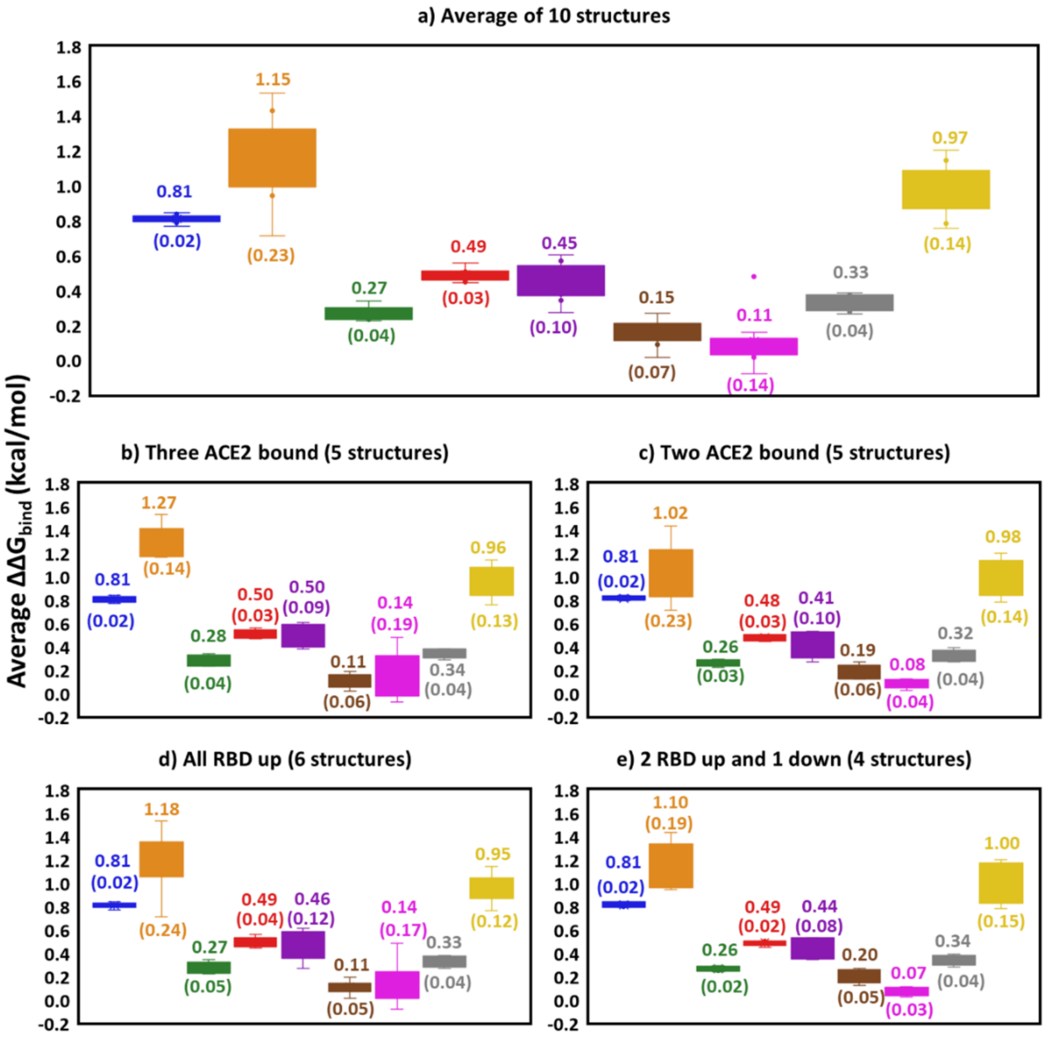


**Figure S7. Average change in binding affinity ($\Delta\Delta G_{bind}$): Comparison of ACE2 complexes with different RBD conformation of S-protein and ACE2 binding. a)** Average $\Delta\Delta G_{bind}$ comparison of 10 ACE2 bound structures for nine mutation categories. The ten ACE2 complexes can be further classified into two categories (**b** and **c**) based on the number of ACE2 bound to S-protein RBD, and into two classes (**d** and **e**) based on the RBD conformation. Average values are shown for each mutation category with standard deviations in brackets.

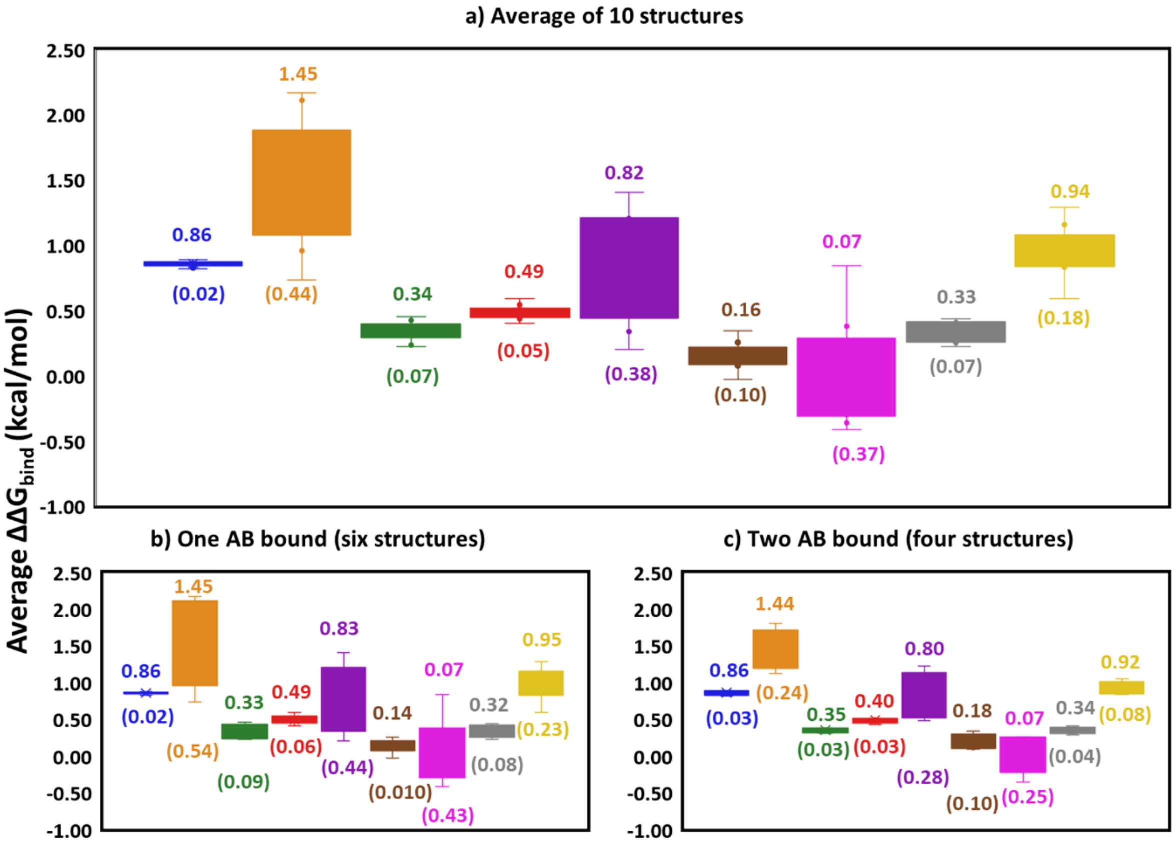


**Figure S8. Average change in binding affinity (ΔΔ$G_{bind}$) comparison of AB complexes with different number of AB units interacting to S-protein RBD region. a)** Average ΔΔ$G_{bind}$ value comparison of 10 AB-bound structures for nine mutation categories. The ten AB complexes can be further classified into two categories (**b** and **c**) based on the number of AB unit bound to S-protein RBD. Average values are shown for each mutation category with standard deviations shown in brackets.

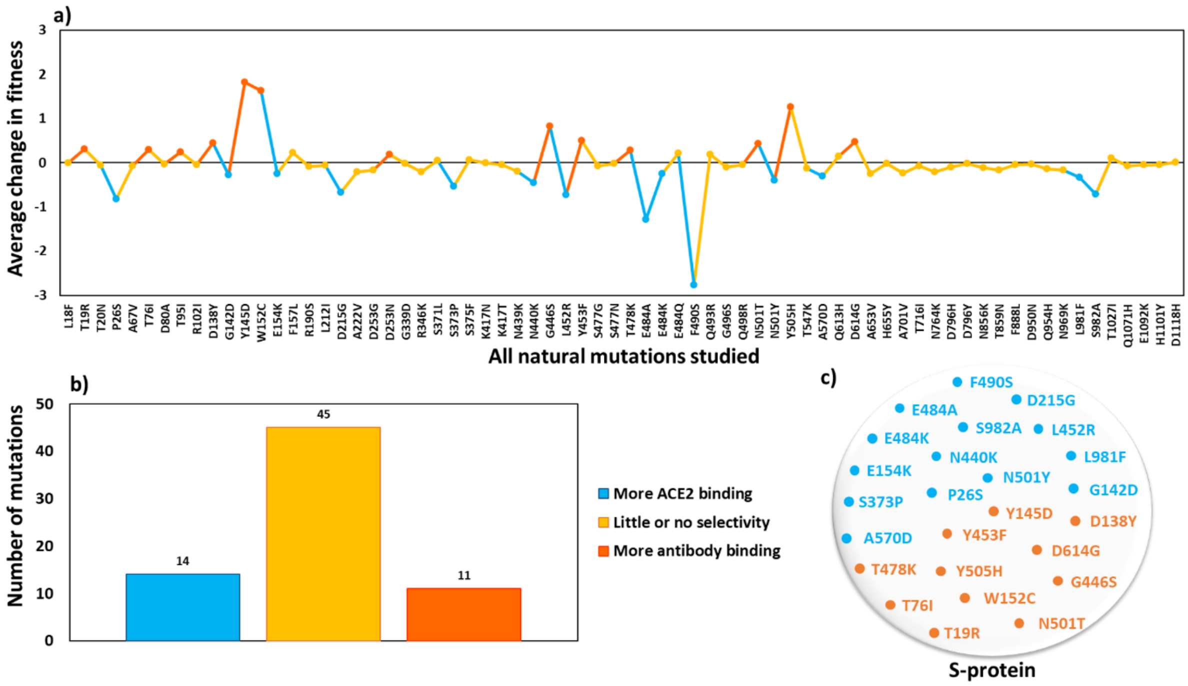


**Figure S9. Average fitness value of natural mutations. (a)** Fitness value of each natural mutation is shown. **(b)** Number of natural mutations showing selective binding towards ACE2/AB/none. **(c)** Naturally evolved sites showing higher binding towards ACE2 or AB are named. For all three plots, mutation showing higher binding towards ACE2, and AB are represented in blue and orange color respectively, whereas mutations with no or little selectivity are shown in yellow colour.

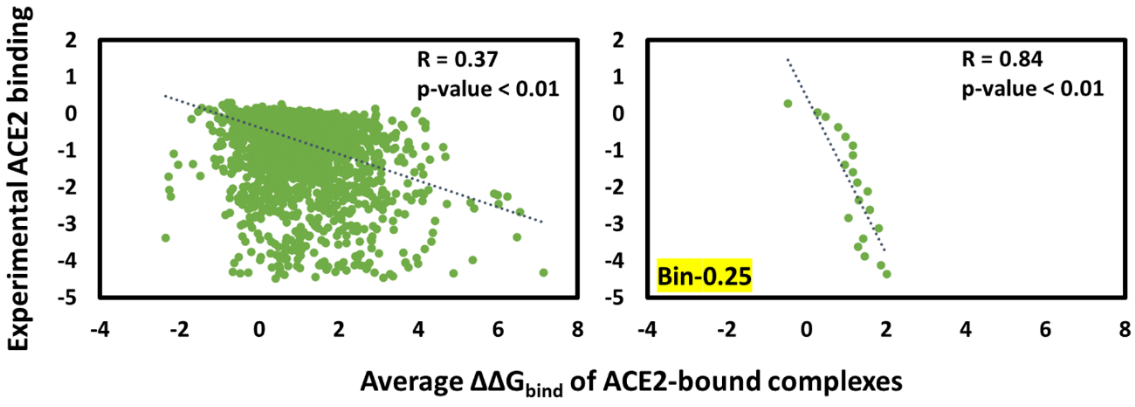


**Figure S10. Scatter plot for comparison of experimental ACE2 binding (Bloom[23] data) with the predicted affinity change (average $\Delta\Delta G_{bind}$) of each mutation in ACE2-bound complexes.** The datapoints present in all the 10 PDB structures and common with the experimental dataset were considered. A total of 3283 data points were plotted.

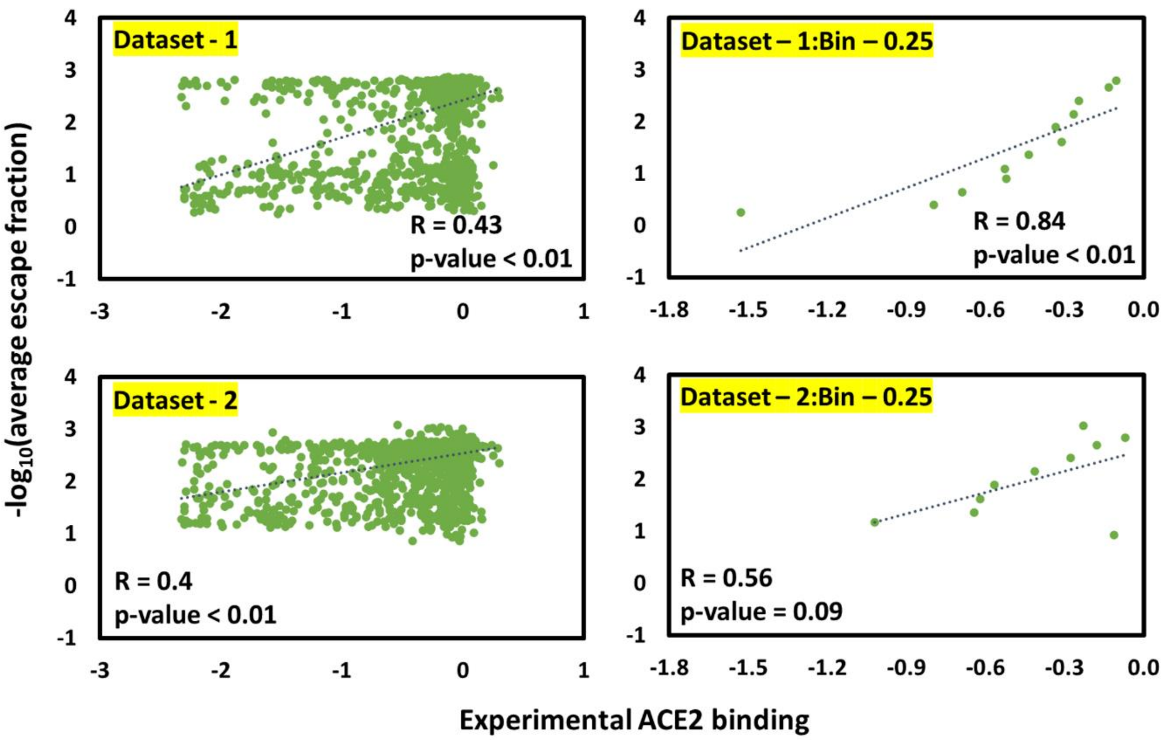


**Figure S11: Scatter plots for comparison of experimental ACE2 binding (dataset 0) with experimental escape fraction values (-$\log_{10}$(average escape fraction); for datasets 1 and 2).** Comparison of dataset 0 with datasets 1 and 2 contain 1915 and 2334 data points respectively. A higher value of ACE2 binding shows a higher ACE2 binding whereas a higher value of -$\log_{10}$(average escape fraction) indicates a high AB binding. Zero average escape fraction values were excluded during negative logarithmic conversion. Correlation increased when bin approach (0.25 bins) was used. The correlation and direction suggest that the S-protein mutations that decrease binding for one protein (ACE2) also decrease for the other protein, in this case the antibodies.

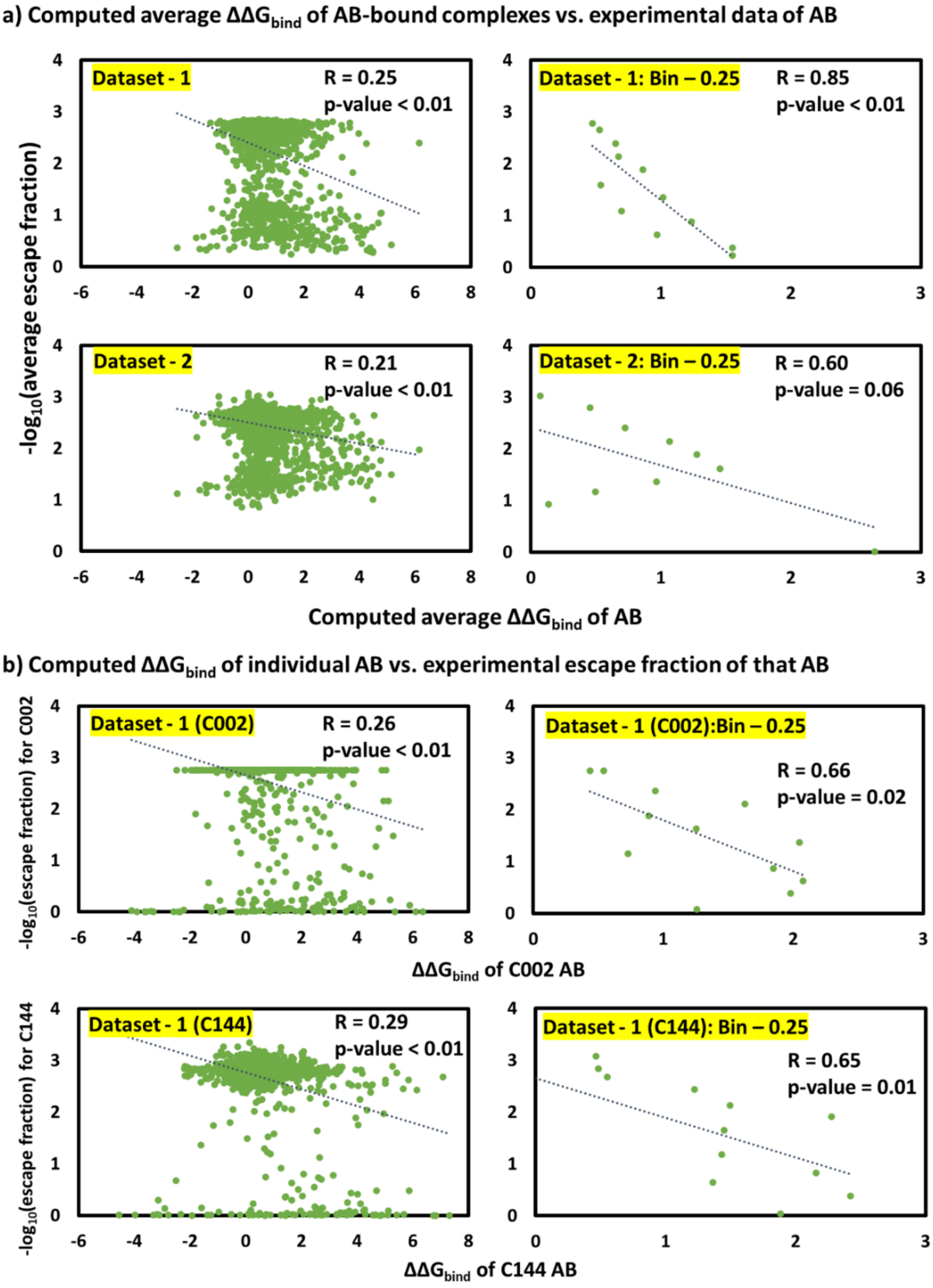


**Figure S12. Comparison of computed and experimental mutation effects for antibodies. a)** Plots show the correlation between computed binding of AB complexes with experimental binding escape data taken from **dataset-1** (1924 datapoints from Greaney et al.[24]) and **dataset-2** (2346 datapoints from Greaney et al.[25]). Zero escape fraction values (nine data points) were found in dataset 2 and excluded from the analysis during negative logarithmic conversion. **b)** Computed $\Delta\Delta G_{bind}$ value of individual AB is correlated with its experimental escape fraction values, which were only available for C002 (PDB ID – 7K8S) and C144 (PDB ID – 7K90) in **dataset-1** (Greaney et al.[24]). 1950 and 1949 datapoints for C002 and C144 were used. One point with zero escape fraction values was excluded from C144 dataset during logarithmic conversion.

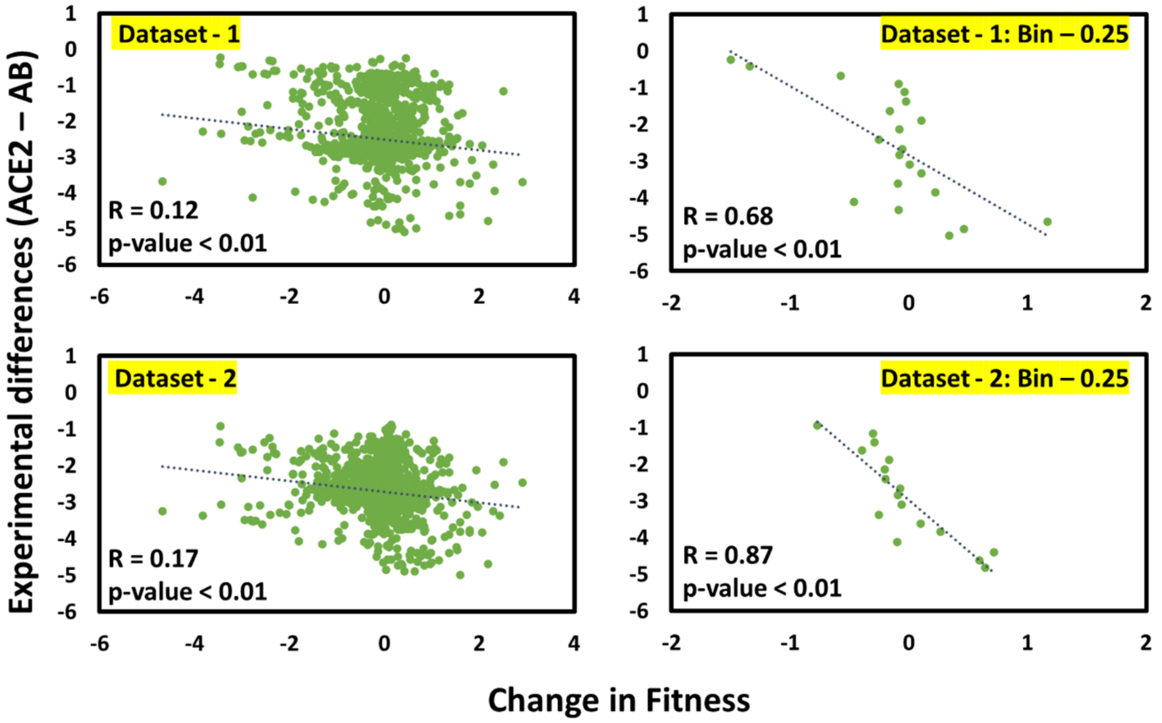


**Figure S13. Experimental vs. computed differential binding effects on ACE2 and antibodies, averaged over all structures for dataset - 1 (1847 points) and dataset – 2 (2245 points).** Change in fitness = Average $\Delta\Delta G_{bind}$ (ACE2 – AB), where a low value indicates a high ACE2 binding. Experimental difference (ACE2 – AB) is the difference of average experimental ACE2 binding and -$\log_{10}$(average AB escape fraction). A high value of experimental difference indicates higher ACE2 binding.

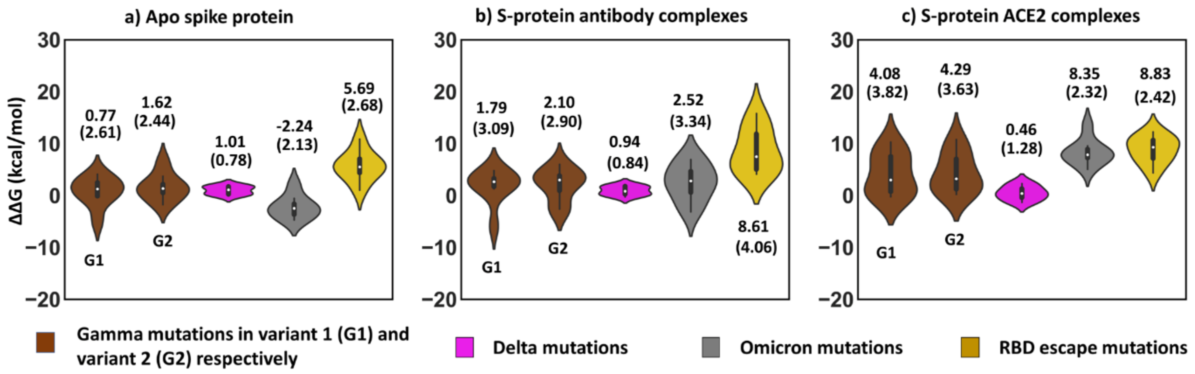


**Figure S14. Epistasis study – ΔΔG values of eight structures of each apo, AB, and ACE2 plotted for four variants (Gamma variant 1, Gamma variant 2, Delta, and Omicron), and RBD escape mutations.** The average value is shown in each plot with standard deviation in brackets. The ΔΔG values were calculated using FoldX[26] by applying all mutations simultaneously to observe the epistasis effect.

[14]"Centers for Disease Control and Prevention," (n.d.).

[15]"WHO," (n.d.).

[16] A. Baum, B.O. Fulton, E. Wloga, R. Copin, K.E. Pascal, V. Russo, S. Giordano, K. Lanza, N. Negron, M. Ni, Y. Wei, G.S. Atwal, A.J. Murphy, N. Stahl, G.D. Yancopoulos, and C.A. Kyratsous, "Antibody cocktail to SARS-CoV-2 spike protein prevents rapid mutational escape seen with individual antibodies," **369**(6506), 1014–1018 (2020).

[17] S.J. Zost, P. Gilchuk, J.B. Case, E. Binshtein, R.E. Chen, J.P. Nkolola, A. Schäfer, J.X. Reidy, A. Trivette, R.S. Nargi, R.E. Sutton, N. Suryadevara, D.R. Martinez, L.E. Williamson, E.C. Chen, T. Jones, S. Day, L. Myers, A.O. Hassan, N.M. Kafai, E.S. Winkler, J.M. Fox, S. Shrihari, B.K. Mueller, J. Meiler, A. Chandrashekar, N.B. Mercado, J.J. Steinhardt, K. Ren, Y.M. Loo, N.L. Kallewaard, B.T. McCune, S.P. Keeler, M.J. Holtzman, D.H. Barouch, L.E. Gralinski, R.S. Baric, L.B. Thackray, M.S. Diamond, R.H. Carnahan, and J.E. Crowe, "Potently neutralizing and protective human antibodies against SARS-CoV-2," Nature **584**(7821), 443–449 (2020).

[18] D.J. Benton, A.G. Wrobel, P. Xu, C. Roustan, S.R. Martin, P.B. Rosenthal, J.J. Skehel, and S.J. Gamblin, "Receptor binding and priming of the spike protein of SARS-CoV-2 for membrane fusion," Nature **588**(7837), 327–330 (2020).

[19] L. Guo, W. Bi, X. Wang, W. Xu, R. Yan, Y. Zhang, K. Zhao, Y. Li, M. Zhang, X. Cai, S. Jiang, Y. Xie, Q. Zhou, L. Lu, and B. Dang, "Engineered trimeric ACE2 binds viral spike protein and locks it in 'Three-up' conformation to potently inhibit SARS-CoV-2 infection," Cell Res. **31**(1), 98–100 (2021).

[20] T. Xiao, J. Lu, J. Zhang, R.I. Johnson, L.G.A. McKay, N. Storm, C.L. Lavine, H. Peng, Y. Cai, S. Rits-Volloch, S. Lu, B.D. Quinlan, M. Farzan, M.S. Seaman, A. Griffiths, and B. Chen, "A trimeric human angiotensin-converting enzyme 2 as an anti-SARS-CoV-2 agent," Nat. Struct. Mol. Biol. **28**(2), 202–209 (2021).

[21] T. Zhou, Y. Tsybovsky, J. Gorman, M. Rapp, G. Cerutti, G.-Y. Chuang, P.S. Katsamba, J.M. Sampson, A. Schön, J. Bimela, J.C. Boyington, A. Nazzari, A.S. Olia, W. Shi, M. Sastry, T. Stephens,

J. Stuckey, I.-T. Teng, P. Wang, S. Wang, B. Zhang, R.A. Friesner, D.D. Ho, J.R. Mascola, L. Shapiro, and P.D. Kwong, "Cryo-EM Structures of SARS-CoV-2 Spike without and with ACE2 Reveal a pH-Dependent Switch to Mediate Endosomal Positioning of Receptor-Binding Domains," Cell Host Microbe **28**(6), 867-879.e5 (2020).
